# Visual to default network pathways: A double dissociation between semantic and spatial cognition

**DOI:** 10.1101/2023.11.28.568861

**Authors:** Tirso RJ Gonzalez Alam, Katya Krieger-Redwood, Dominika Varga, Zhiyao Gao, Aidan Horner, Tom Hartley, Michel Thiebaut de Schotten, Magdalena W Sliwinska, David Pitcher, Daniel S. Margulies, Jonathan Smallwood, Elizabeth Jefferies

## Abstract

Processing pathways between sensory and default mode network (DMN) regions support recognition, navigation, and memory but their organisation is not well understood. We show that functional subdivisions of visual cortex and DMN sit at opposing ends of parallel streams of information processing that support visually-mediated semantic and spatial cognition, providing convergent evidence from univariate and multivariate task responses, intrinsic functional and structural connectivity. Participants learned virtual environments consisting of buildings populated with objects, drawn from either a single semantic category or multiple categories. Later, they made semantic and spatial context decisions about these objects and buildings during functional magnetic resonance imaging. A lateral ventral occipital to frontotemporal DMN pathway was primarily engaged by semantic judgements, while a medial visual to medial temporal DMN pathway supported spatial context judgements. These pathways had distinctive locations in functional connectivity space: the semantic pathway was both further from unimodal systems and more balanced between visual and auditory-motor regions compared with the spatial pathway. When semantic and spatial context information could be integrated (in buildings containing objects from a single category), regions at the intersection of these pathways responded, suggesting that parallel processing streams interact at multiple levels of the cortical hierarchy to produce coherent memory-guided cognition.

## Introduction

The Default Mode Network (DMN) is involved in higher-order cognition including in semantic cognition, mental time travel and scene construction (Andrews-Hanna et al., 2010b, 2010a; Lambon Ralph et al., 2017; Spreng et al., 2009). Its functions and architecture are plagued by apparent contradictions: it often deactivates in response to visual inputs yet it is connected to visual cortex (Knapen, 2021; Leech et al., 2012; Szinte and Knapen, 2020). In addition, this network is associated with both abstraction from sensory-motor features (Chiou et al., 2020a, 2019; Gonzalez Alam et al., 2021; Rice et al., 2015b, 2015a) and internally-generated states like imagery and autobiographical memory (Philippi et al., 2015; Ritchey and Cooper, 2020; Spreng and Grady, 2010; Zhang et al., 2022). A recent perspective suggests these diverse functions are facilitated by the topographical location of DMN on the cortical mantle (Smallwood et al., 2021). DMN is maximally separated from sensory-motor regions – both in terms of its physical location and in connectivity space. It is at one end of the principal gradient of intrinsic connectivity that captures the separation of unimodal and heteromodal cortex (Margulies et al., 2016) and this location is thought to allow DMN to sustain representations that are distinct from sensory-motor features and at odds with the current state of the external world (Murphy et al., 2019, 2018).

Despite these common functional characteristics of DMN, parcellations of intrinsic connectivity reveal subdivisions (Andrews-Hanna et al., 2010b; Schaefer et al., 2018; Wen et al., 2020; Yeo et al., 2011). Lateral fronto-temporal (FT) DMN regions are associated with semantic cognition, including the abstraction of heteromodal meanings from sensory-motor features and the ability to access these meanings from sensory inputs in a task-appropriate way (Chiou et al., 2020a, 2019; Lambon Ralph et al., 2017; Wang et al., 2020). In contrast, scene construction, thought to be a key component of episodic recollection, is associated with a medial temporal (MT) subsystem (Andrews-Hanna et al., 2010b; D’Argembeau et al., 2010; Hassabis et al., 2007; Zhang et al., 2022). FT and MT-DMN subnetworks are interdigitated in regions of core DMN (Braga and Buckner, 2017) and they are assumed to work together but little is known about how information within them is integrated. One hypothesis is that the spatial adjacency of DMN subsystems allows their common recruitment and coordination when semantic and scene-based information is aligned; for example, when semantically similar objects are found in a common location, or spatial position predicts the meanings of items that are found there.

FT and MT-DMN support heteromodal representations and yet can be accessed by visual inputs, raising the question of how neural pathways between vision and DMN are organised. Visual neuroscience has revealed different responses associated with recognising objects and scenes (Kravitz et al., 2013, 2011). Objects engage a ventral pathway extending laterally and anteriorly through ventral lateral occipital cortex (LOC) and the fusiform gyrus towards the anterior temporal lobes (ATL), thought to be a key heteromodal hub for conceptual representation. This pathway might act as input to FT-DMN (Andrews-Hanna et al., 2014a; Andrews-Hanna and Grilli, 2021a; DiCarlo et al., 2012; Kravitz et al., 2013; Malach et al., 2002). Navigating visuospatial environments and scene construction, on the other hand, involves the occipital place area, posterior cingulate, retrosplenial, entorhinal and parahippocampal cortex, before this pathway terminates in hippocampus. These regions are associated with the MT-DMN subnetwork (Andrews-Hanna et al., 2014a; Andrews-Hanna and Grilli, 2021a; Epstein and Baker, 2019; Kravitz et al., 2011; Reagh and Yassa, 2014). This work suggests that visual and DMN subsystems may be linked. For example, during memory for people and places, medial parietal cortex mirrors the well-established medial-lateral organisation of ventral temporal cortex during the perception of scenes and faces; medial parietal regions also show differential connectivity to these visual regions (Margulies et al., 2009; Silson et al., 2019; Steel et al., 2021). Yet these past studies did not examine whole-brain connectivity or semantic cognition beyond the social domain and were also unable to explore the interaction of these pathways.

Here, we used multiple neuroscientific methods to delineate the pathways from visual cortex to DMN, providing convergent evidence for two parallel streams supporting semantic and spatial cognition. In Study 1, participants learned about virtual environments (buildings) populated with objects belonging to diverse semantic categories, both man-made (tools, musical instruments, sports equipment) and natural (land animals, marine animals, birds). We then used fMRI to examine neural activity as participants viewed object and scene probes and made semantic and spatial context decisions. Some buildings were associated with a specific semantic category (e.g., a building filled with musical instruments), while others included a mix of categories, allowing us to examine the interaction between semantic and spatial cognition. We identified dissociable pathways of connectivity between different parts of visual cortex and DMN subsystems; these overlapped with visual localiser responses for objects and scenes (in Study 2), as well as previously described DMN subsystems, and showed different patterns of functional and structural connectivity (in Study 3). These pathways refer to regions that are coupled, functionally or structurally, together, providing the potential for communication between them. They also had distinctive locations in a functional space defined using whole-brain gradients of connectivity: the semantic pathway was further from unimodal systems and more balanced between visual and auditory-motor regions compared with the spatial pathway. Moreover, when semantic and spatial context information could be combined (e.g. when the objects in a building were from the same semantic category), regions at the intersection of these pathways responded, in both DMN and visual cortex, suggesting these parallel processing streams can interact at multiple levels of the cortical hierarchy to produce coherent memory-guided cognition.

## Results

### Behavioural Results

To examine task accuracy, we performed a 2×2 repeated-measures ANOVA using task (2 levels: semantic, spatial context) and condition (2 levels: Mixed-Category Building (MCB), and Same-Category Building (SCB)) as factors. There was a main effect of task (F(1,26)=76.52, p<.001), condition (F(1,26)=11.31, p=.002) and a task by condition interaction (F(1,26)=14.51, p<.001). Participants showed poorer accuracy in the spatial context task relative to the semantic task and in the MCB relative to the SCB condition. Participants were significantly less accurate in the MCB trials relative to the SCB trials of the spatial context task (t(26)=4.08, p<.001); this difference was not observed in the semantic task (t(26)=.74, p=.47).

Response times showed the same pattern, with main effects of task (F(1,26)=51.37, p<.001), condition (F(1,26)=31.14, p<.001) and their interaction (F(1,26)=29.48, p<.001). Participants had slower reaction times in the spatial context task than the semantic task and in the MCB relative to the SCB condition. Post-hoc comparisons confirmed that participants were significantly slower in MCB than SCB trials of the spatial context task (t(26)=6.08, p<.001), but this difference was not observed in the semantic task (t(26)=.1, p=.92). These results are shown in Figure 1.

**Figure 1.**
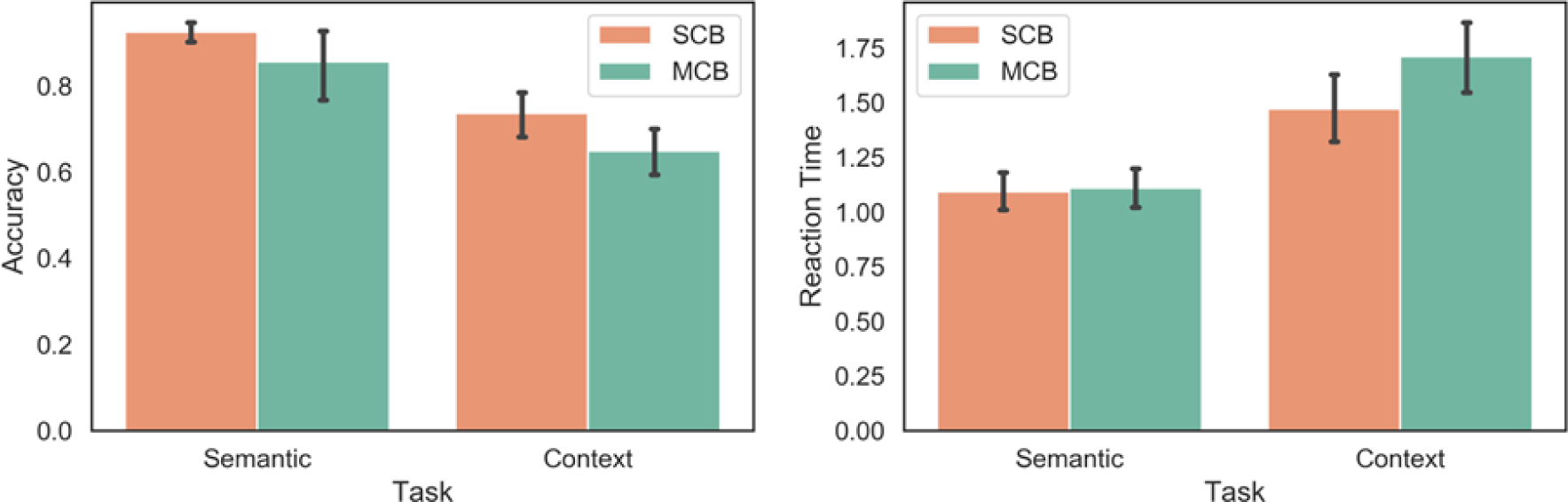
Behavioural results for the semantic and spatial context tasks inside the scanner. SCB = Same Category Buildings: all the items in the building were taken from the same semantic category. MCB = Mixed Category Buildings: the items in the buildings were drawn from different semantic categories.

### Neuroimaging Results

To probe the organisation of streams of information between visual cortex and DMN, our neuroimaging analysis strategy consisted of a combination of task-based and connectivity approaches. We first delineated the regions in visual cortex that are engaged by the viewing of probes during our task (Figure 2), as well as the DMN regions that respond when making decisions about those probes (Figure 3): we characterised both by comparing the activation maps with well-established DMN and object/scene perception regions, analysed the pattern of activation within them, their functional connectivity and task associations. Having characterised these dissociable visual and DMN regions, we proceeded to ask whether they are differentially linked: are the visual regions activated by object probe perception more strongly linked to DMN regions that are activated when making semantic decisions about object probes, relative to other DMN regions? Is the same true for visual regions associated with scene perception and DMN regions responding to spatial decisions about which rooms were in the same building? We answered this question through a series of connectivity analyses (Figure 4) that examined: 1) if the functional connectivity of visual to DMN regions (and DMN to visual regions) shows a dissociation, suggesting there are object semantic and spatial cognition processing ‘pathways’; 2) if this pattern was replicated in structural connectivity; 3) if it was present at the level of individual participants, and, 4) we characterised the spatial layout, network composition (using influential RS networks) and cognitive decoding of these pathways. Having found dissociable pathways for semantic (object) and spatial context (scene) processing, we then examined their position in a high-dimensional connectivity space (Figure 5) that allowed us to document that the semantic pathway is less reliant on unimodal regions (i.e., more abstract) while the spatial context pathway is more allied to the visual system. Finally, we used uni- and multivariate approaches to examine how integration between these pathways takes place when semantic and spatial information is aligned (Figure 6).

**Figure 2:**
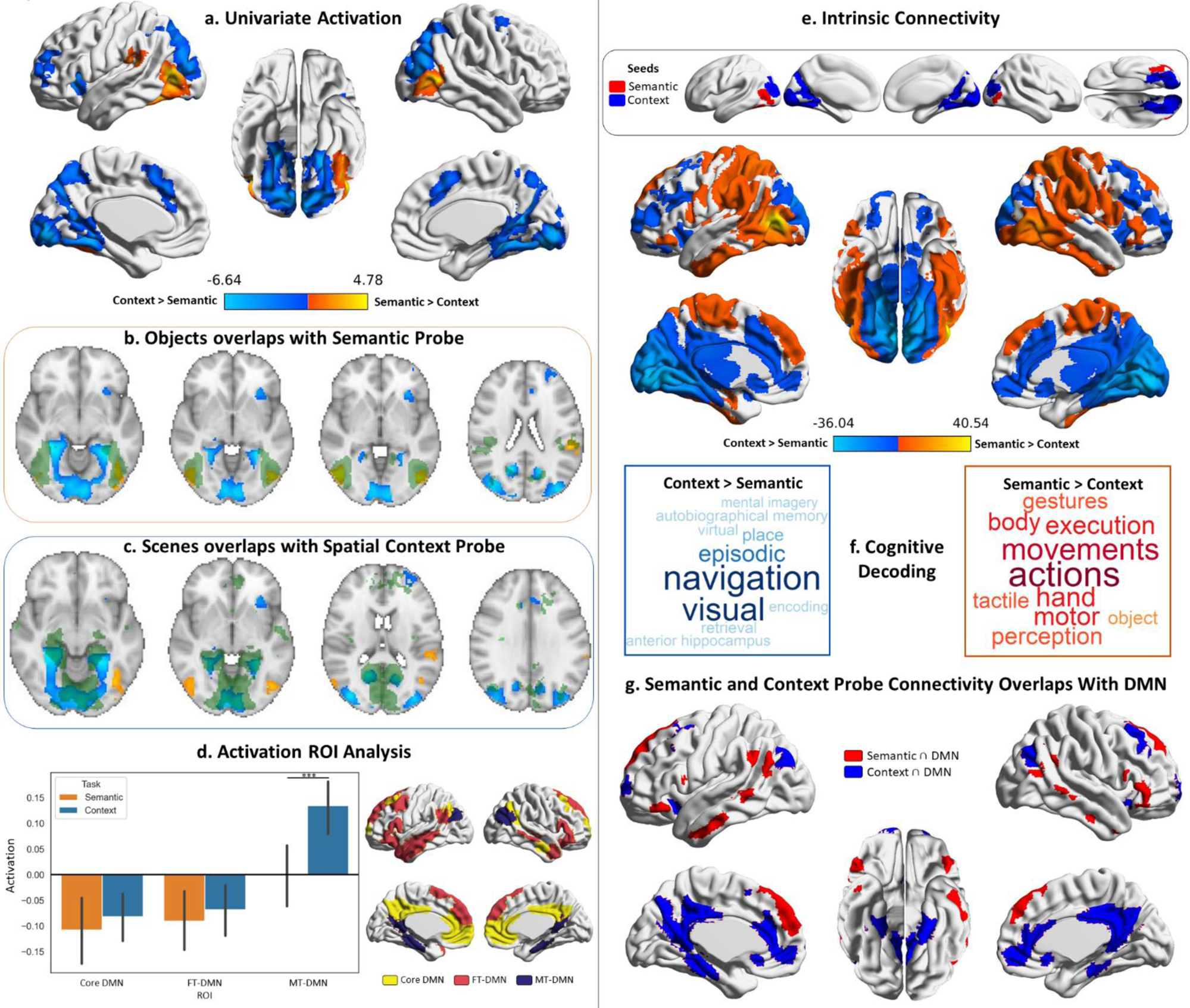
*Probe responses*. Warm colours = semantic > spatial context probes. Cool colours = spatial context > semantic probes. Left panel: Univariate results from Study 1, contrasting semantic and spatial context probes. Right panel: Intrinsic connectivity results from Study 3 using semantic and spatial context probe activation within visual networks as seeds. Panel a: Brain maps depicting the supra-threshold univariate activation results for the probe phase of the semantic and spatial context tasks. Panels b and c: Axial slices showing the overlap of these univariate results with Scene and Object localiser maps from Study 2 (the localiser maps are in green, and the univariate results maps are in warm and cool colours). Panel d: ROI analysis examining the activation in the three default mode subnetworks of the Yeo 17 parcellation during the probe phase of the semantic and spatial context tasks. The error bars in the bar plots depict the standard error of the mean (Note: ***p<.001); the ROIs are shown to the right of the bar plots. Panel e: Brain maps depicting the seeds and intrinsic connectivity results for the semantic and spatial context probe regions. Panel f: Word clouds depicting the cognitive decoding of unthresholded connectivity maps for semantic and spatial context probe seeds using Neurosynth (bigger words reflect stronger correlation of the functional maps with the terms); the colour-code follows that of the brain maps. Panel g: Brain maps showing the overlap of these intrinsic connectivity maps for semantic and spatial context probes with the default mode network from the 7-network parcellation from Yeo et al. (2011).

**Figure 3:**
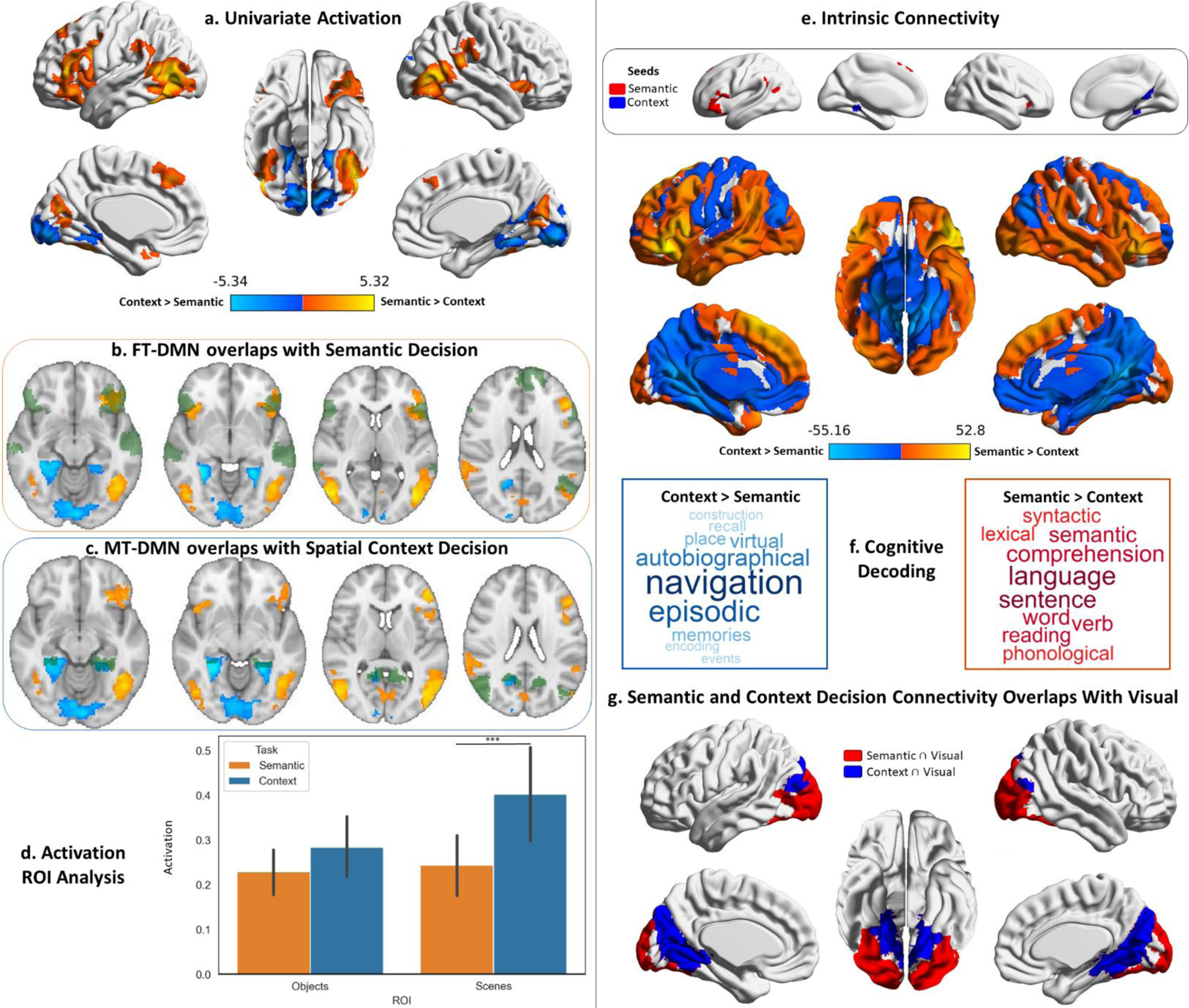
*Decision responses*. Warm colours = semantic > spatial context decisions. Cool colours = spatial context > semantic decisions. Left panel: Univariate results from Study 1 contrasting semantic and spatial context decisions. Right panel: Intrinsic connectivity results from Study 3 using semantic and spatial context decision activation within DMN networks as seeds. Panel a: Brain maps depicting the supra-threshold univariate activation results for the decision phase of the semantic and spatial context tasks. Panels b and c: Axial slices showing the overlap of these univariate results with the FT and MT default mode subnetworks of the Yeo 17 network parcellation (the default mode maps are in green, and the univariate results maps are in warm and cool colours). Panel d: ROI analysis examining the activation in the Scene and Object localiser maps from Study 2 during the decision phase of the semantic and spatial context tasks. The error bars in the bar plots depict the standard error of the mean (Note: ***p<.001, * p < .05); the ROIs are shown in Supplementary Figure S3. Panel e: Brain maps depicting the seeds and intrinsic connectivity results for the semantic and spatial context decision regions. Panel f: Word clouds depicting the cognitive decoding of unthresholded connectivity maps for semantic and spatial context decision seeds using Neurosynth (bigger words reflect stronger correlation of the functional maps with the terms); the colour-code follows that of the brain maps. Panel g: Brain maps showing the overlap of these intrinsic connectivity maps with the visual network from the 7-network parcellation from Yeo et al. (2011).

**Figure 4.**
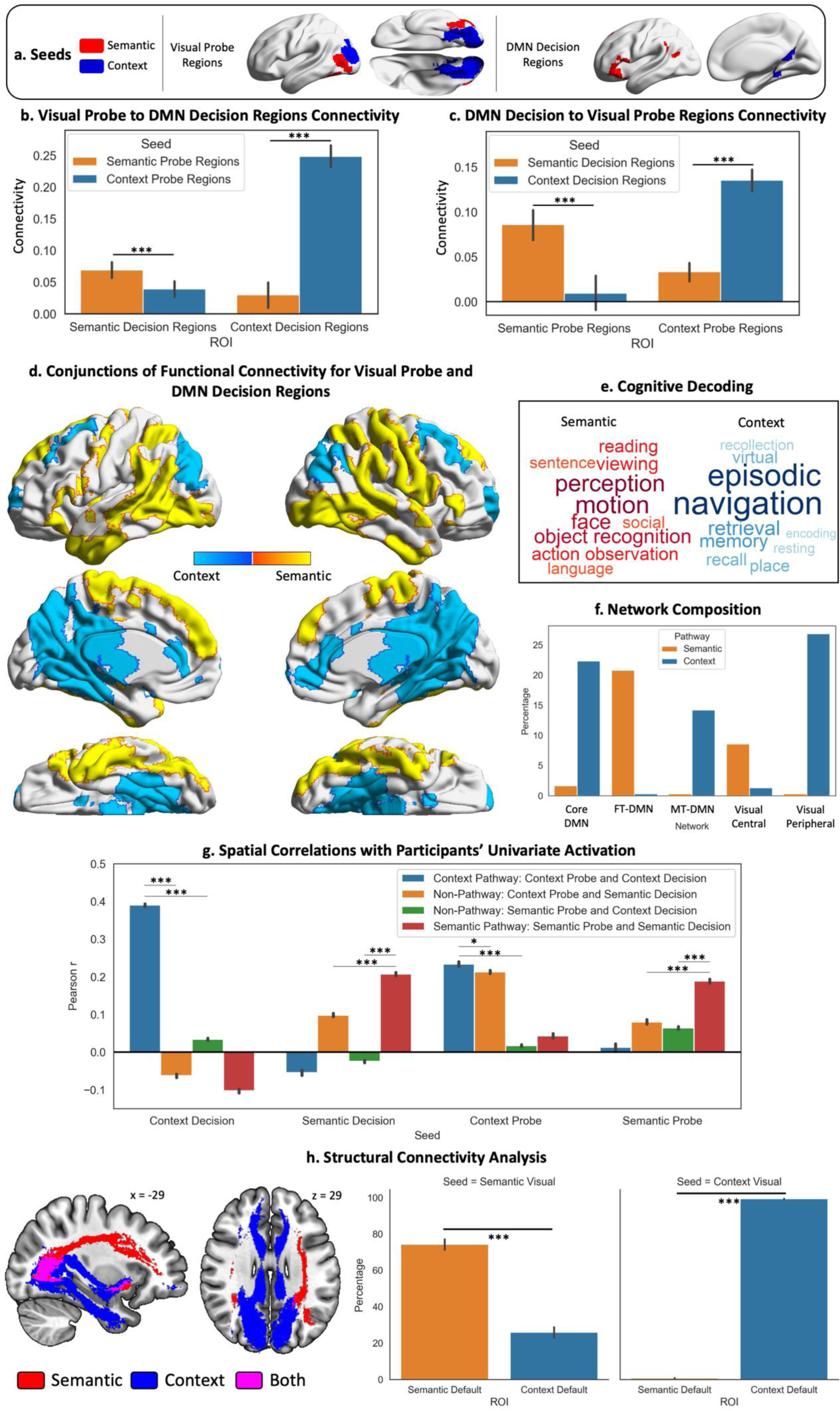
*Panels a, b and c:* These panels depict the seeds, ROIs and their connectivity. The bar plots in panels b and c show the connectivity between DMN decision regions and probe visual regions. *Panel d*: Warm colours = common regions showing stronger intrinsic connectivity to semantic decision regions in DMN and semantic probe regions in visual cortex; Cool colours = common regions showing stronger intrinsic connectivity to spatial context decision regions in DMN and spatial context probe regions in visual cortex. *Panel e:* The cognitive decoding of these spatial maps using Neurosynth following the same colour code as Panel d. *Panel f*: Network composition showing the percentage of each pathway map overlapping with the three DMN and two Visual subnetworks defined by the Yeo et al. (2011) 17-network parcellation. *Panel g:* Results of spatial correlation analysis comparing the semantic and spatial context pathways with non-pathway maps (derived from the conjunction of the connectivity of probe and decision seeds across different tasks, e.g., probe spatial context ∩ decision semantic connectivity). We assessed the spatial similarity of these pathway and non-pathway maps to the univariate activation during the probe and decision phases for each task and each participant. *Panel h:* Results of the structural connectivity analysis. Tracts displayed are a conjunction of streamlines between the probe and decision seeds of each task. The y axis of the bar plots shows the percentage of streamlines from each visual seed that terminate in each DMN ROI (shown in the x axis). The error bars depict the standard error of the mean. *** p < .001, * p < .05.

**Figure 5.**
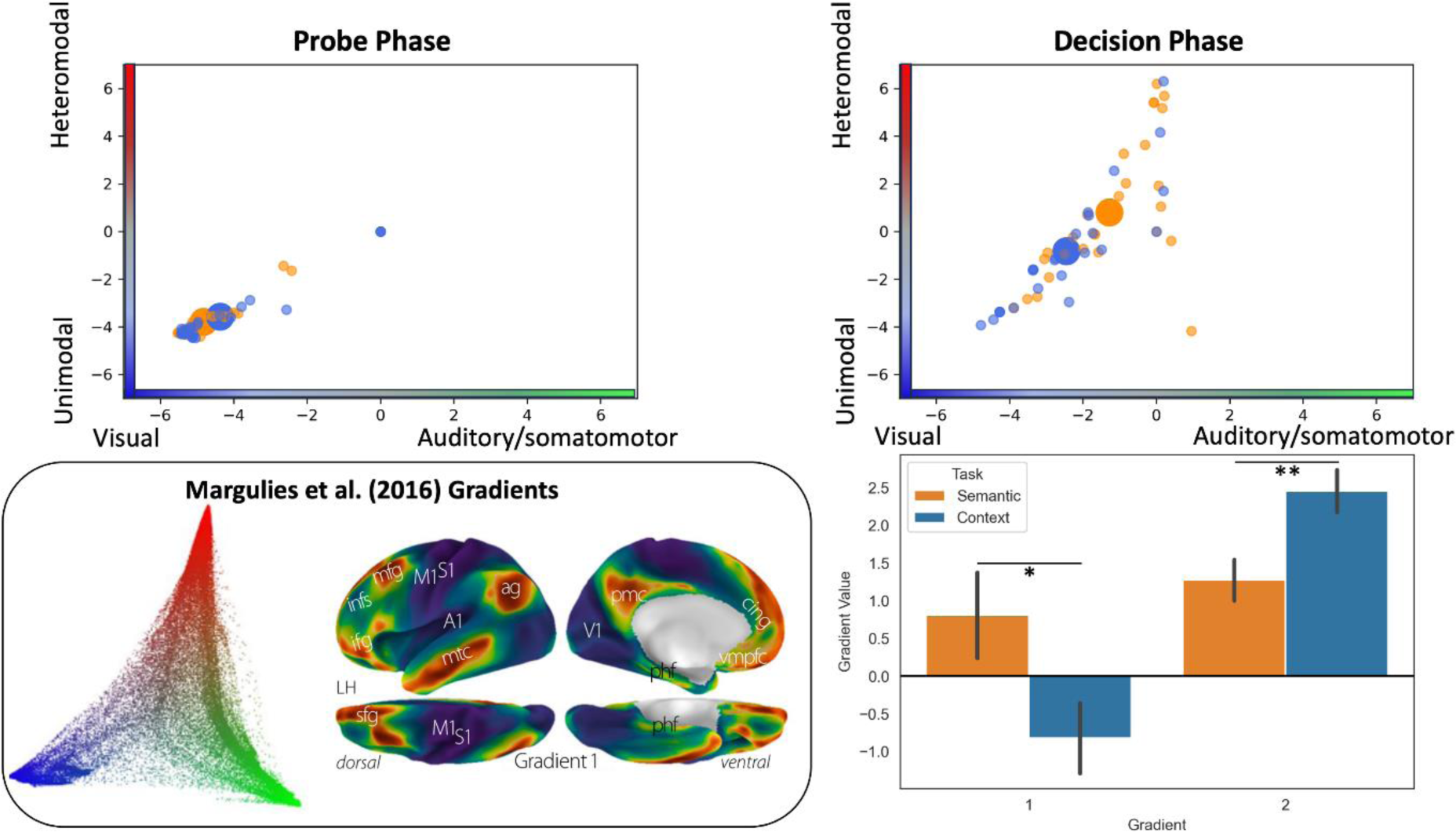
Analysis situating the position of the pathways in a whole-brain connectivity gradient space (Margulies et al., 2016). The scatterplots depict the position of each participant’s peak response to the semantic and context task in this gradient space (the big circles represent the mean of each task for that phase). The bar plots compare the mean of each gradient across tasks. The inset on the bottom left of the panel displays Margulies’ et al. (2016) original gradient space.

**Figure 6.**
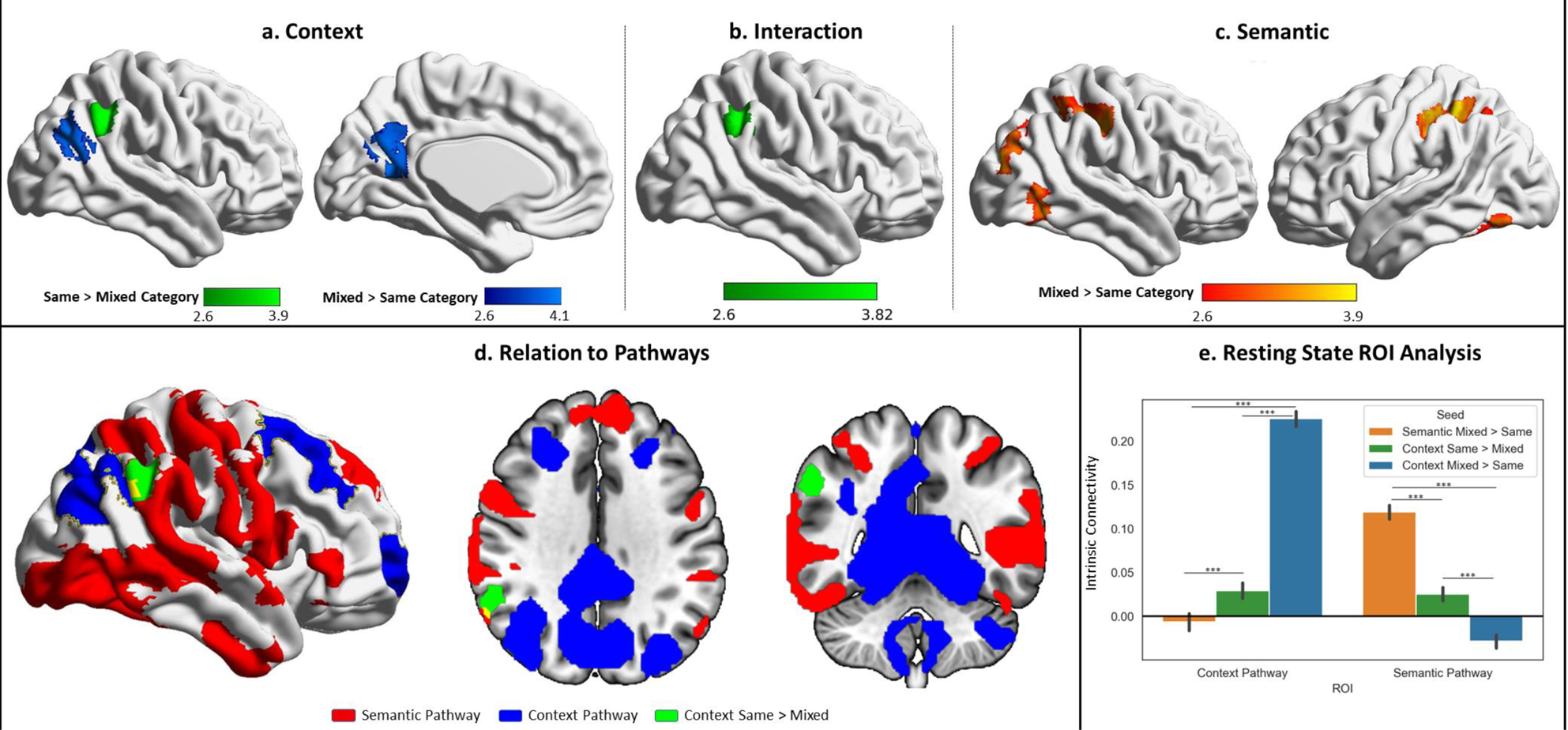
Univariate results for the probe phase contrasting same category vs. mixed category trials separately for the semantic and spatial context tasks. Panel a: Contrast of spatial context same > mixed-category building trials during the probe phase. Panel b: Task by condition (same/mixed-category building) interaction. Panel c: Contrast of semantic same > mixed-category building trials during the probe phase. Panel d: Spatial relations of the same > mixed spatial context cluster with the semantic and spatial context pathways outlined in Figure 4. Panel e: Intrinsic connectivity seed-to-ROI results using the three univariate results clusters shown in the top panel as seeds and the pathways as ROIs. The error bars depict the standard error of the mean. *** p<.001.

#### Probe phase

We began our exploration of the streams of information between visual cortex and DMN by characterising their visual end. To accomplish this, we first analysed the whole-brain activation observed during the probe phase of our semantic and spatial context tasks, when participants were viewing objects and scenes, and related these responses to previously established visual regions for object and scene perception. We then linked the visual regions engaged by our task to the DMN by describing their patterns of intrinsic connectivity, their functional involvement and activation found within DMN regions during the probe phase of our task.

We examined differences in neural responses to probe images of objects in the semantic task, and scenes in the spatial context task in Study 1 (Figure 2). Semantic probes elicited greater activation in bilateral ventral lateral occipital cortex, extending to fusiform cortex and supramarginal gyrus in the left hemisphere. Spatial context probes elicited a stronger response in bilateral dorsal lateral occipital cortex, medial occipital lobe, precuneus, parahippocampal cortex and supplementary motor areas, as well as insula and middle frontal gyrus and frontal pole regions in the left hemisphere, and precentral regions in the right hemisphere (Figure 2, panel a). Details of peak activations for all univariate results from Study 1 are in Supplementary Table S1.

To confirm these distinctive responses to semantic and spatial context probes were related to well-established categorical effects within visual cortex, we examined their overlap with object and scene localisers (i.e. passive viewing) in Study 2 (Figure 2, panels b and c). Regions engaged by the spatial context probes resembled scene perception regions, while semantic probes overlapped with object perception regions.

We next analysed the probe responses exploring the strength of activation during this phase within three DMN subdivisions defined by Yeo et al. (2011)^1^. A two-way repeated measures ANOVA using task (semantic, spatial context) and ROI (Core DMN, FT-DMN, MT-DMN) as factors revealed a significant main effect of ROI (F(1.281,33.306)=50.42, p<.001) and an interaction (F(1.64,42.65)=12.44, p<.001; Greenhouse-Geisser corrected). Post-hoc comparisons showed a significantly stronger response within MT-DMN to spatial context relative to semantic probes (t(26)=4.1, p=.001). No significant difference between tasks was observed for core or FT-DMN (both p>.05, Figure 2, panel d).

Finally, we examined the intrinsic connectivity of activation regions in Figure 2a, masked by Yeo et al.’s (2011) visual networks (combining central and peripheral networks), using data from Study 3. Visual areas responding to semantic and spatial context probes showed differential connectivity, including to regions of DMN (posterior cingulate, medial prefrontal cortex, portions of anterior and dorsal prefrontal cortex and anterior temporal cortex; Figure 2, panel e). Cognitive decoding using Neurosynth revealed that semantic probe connectivity was associated with perceptual and somatomotor terms, while spatial context probe connectivity was associated with navigation, visuospatial and episodic memory terms (Figure 2, panel f). Semantic probe regions showed preferential overlap with FT-DMN, whilst spatial context probe regions showed greater overlap with MT-DMN, followed by core DMN (Figure 2, panel g).

#### Decision phase

Having characterised the visual end of the visual-DMN pathways in our previous analysis, we next turned our attention to the DMN end. Following the same logic, we first analysed the whole-brain activation during the decision phase of our task, where participants had to judge the relationship between objects and scenes respectively. We compared this activation to classic DMN regions described by Yeo et al. (2011) and linked the regions engaged within this network to visual regions in terms of their task activation and functional connectivity.

We characterised regions responsive to semantic and spatial context decisions in Study 1. Figure 3a, shows that semantic decisions elicited stronger engagement within dorsolateral prefrontal, lateral occipital, posterior temporal and occipital cortex, as well as pre-SMA. These regions overlapped with FT-DMN (Figure 3, panel b). Spatial context decisions produced stronger activation within a predominantly medial set of occipital, ventromedial temporal (including parahippocampal gyrus), retrosplenial and precuneus regions that overlapped with MT-DMN (Figure 3, panel c).

Given our hypothesis of dissociable pathways between visual cortex and DMN subsystems, we examined responses to semantic and spatial contextual decisions in visual cortex, using the object and scene localiser ROIs from Study 2. A two-way repeated-measures ANOVA including task (semantic, spatial context decisions) and visual ROI (scene and object regions) revealed main effects of task (F(1,26)=14.02, p<.001), visual ROI (F(1,26)=9.91, p=.004) and their interaction (F(1,26)=24.65, p<.001). Post-hoc comparisons showed a stronger response to spatial context decisions, relative to semantic decisions, in the visual ROI sensitive to scenes (t(26)=3.63, p<.001), but not in the object-selective region (t(26)=1.88 p=.071; Figure 3, panel d).

We investigated differences in the intrinsic connectivity of the distinct DMN decision regions activated during the semantic and spatial context tasks (shown in Figure 3a) in independent data from Study 3. We seeded the decision regions that intersected with DMN (combining core, FT and MT-DMN from Yeo et al.’s 2011 parcellation). The results, in Figure 3e, revealed differences in the functional networks of these DMN regions that extended to visual cortex. Semantic decision regions showed stronger connectivity to lateral visual regions along with lateral temporal cortex, inferior frontal gyrus, angular gyrus and dorsomedial prefrontal cortex. Spatial decision regions were more connected to medial visual regions, ventro-medial temporal regions, medial parietal cortex, ventral parts of medial prefrontal cortex, motor cortex and dorsal parts of lateral occipital cortex. Since resting-state analysis is sensitive to the choice of threshold, we repeated this analysis with a stricter cluster-forming threshold and found that the resulting maps were virtually identical (with a spatial correlation of r=.99 between maps generated with both thresholds), for both the probe and decision phases (see Supplementary Materials: Resting-state maps with stricter thresholding).

Cognitive decoding of these connectivity maps using Neurosynth (Figure 3, panel f) revealed that the semantic decision network was associated with terms related to language, semantic processing and reading, while the spatial context decision network was associated with navigation, episodic and autobiographical memory. The decision DMN seeds also showed differential connectivity to visual regions (Figure 3, panel g). Semantic decision regions were more connected with lateral and ventral occipital cortex, whilst spatial context decision regions showed more connectivity with medial occipital, ventromedial temporal (including parahippocampal) and dorsal lateral occipital cortex.

#### Pathways analysis

The analysis above identified regions of visual cortex showing a differential response to semantic and spatial context probes, related to category effects for objects versus scenes. We also found distinct DMN subnetworks which supported semantic and spatial context decisions respectively. Next, we considered if these effects are linked: Do FT-DMN regions have stronger connectivity to object perception areas of visual cortex, while MT-DMN regions connect to scene perception regions? To answer this question, we analysed the functional and structural connectivity from the visual regions that were responsive to viewing probes during our tasks to those DMN regions activated during decisions about those probes. We characterised the spatial layout of these pathways in the brain as well as their large-scale network composition, their functional involvement, and verified their presence in individual participants.

Using resting-state data from Study 3, we performed a series of Seed-to-ROI analyses to examine differential visual to DMN connectivity. We seeded the visual regions in Figure 2e (i.e., probe responses to objects and scenes masked by Yeo et al.’s visual networks) and extracted their intrinsic connectivity to DMN, using ROIs showing differential activation to semantic and spatial context decisions (corresponding to the seeds in Figure 3, panel e). In a second analysis, we examined the reverse (i.e., seeded DMN regions and extracted their connectivity to visual regions). The seeds and ROIs for this analysis can be consulted in Figure 4a. These effects were analysed using repeated-measures ANOVAs examining the interaction between seed and ROI (Figure 4, panels b and c). The Visual-to-DMN ANOVA showed main effects of seed (F(1,190)=226.23, p<.001), ROI (F(1,190)=85.21, p<.001) and a seed by ROI interaction (F(1,190)=322.83, p<.001). Post-hoc contrasts confirmed there was stronger connectivity between object probe regions and semantic versus spatial context decision regions (t(190)=3.98, p<.001), and between scene probe regions and spatial context versus semantic decision regions (t(190)=20.07, p<.001). The DMN-to-Visual ANOVA confirmed this pattern: again, there was a main effect of ROI (F(1,190)=36.91, p<.001) and a seed by ROI interaction (F(1,190)=218.42, p<.001), with post-hoc contrasts confirming stronger intrinsic connectivity between DMN regions implicated in semantic decisions and object probe regions (t(190)=11.63, p<.001), and between DMN regions engaged by spatial context decisions and scene probe regions (t(190)=6.17, p<.001). To ensure that these results were not artificially inflated due to spatial mixing of the resting-state signals arising from proximal visual peripheral and DMN-C networks (Silson et al., 2019; Steel et al., 2021), we conducted a supplementary analysis eroding the visual Probe and DMN Decision ROIs for the spatial context task until the minimum gap between them exceeded the size of our smoothing kernel (see Supplementary Materials: Eroded Masks Replication Analysis and Figure S4). The results replicated the pattern described above. Supplementary analyses using the same seeds and task-independent ROIs also revealed the same pattern: these ROIs were based on the visual localiser masks from Study 2 and the complete DMN subnetworks defined by the Yeo et al. (2011) 17-network parcellation (see Supplementary Analysis: “Replicating resting-state connectivity pathways with task-independent ROIs” and Figure S5).

These pathways, specialised for semantic and spatial cognition, link dissociable visual regions to DMN subsystems, consistent with the suggestion that functional differentiation in DMN partly reflects the strength of different inputs. Figure 4d provides a visualisation of these pathways using the intersection of connectivity from object over scene probe regions and semantic versus spatial context decisions to identify the semantic pathway (warm colours) and the reverse contrasts for the spatial pathway (cool colours). Cognitive decoding revealed terms related to object, action, motion, social and face perception, as well as language and reading for the semantic pathway, and terms related to navigation, place processing and memory for the spatial context pathway (Figure 4, panel e). The semantic pathway was predominantly characterised by FT-DMN and Visual Central regions in the Yeo et al. (2011) 17-network parcellation, whilst the spatial context pathway reflected Core and MT-DMN, and Visual Peripheral networks (Figure 4, panel f).

A complementary analysis examined the spatial correlation of these semantic and spatial pathways (Figure 4d) with participants’ univariate activation when viewing semantic and spatial context probes, and when making semantic and spatial context decisions. We compared spatial correlations between our hypothesised pathways and ‘non-pathway conjunctions’, defined as conjunctions of visual object probe and DMN spatial context decision connectivity, and visual scene probe and DMN semantic decision connectivity with these univariate activation maps. We obtained Pearson r values for each participant reflecting spatial similarity of their activation patterns with these pathway and non-pathway maps and compared these correlations using one-way ANOVAs (Figure 4, panel g). There was a significant effect of pathway for each of the four task phases (Spatial Context Decision: F(2.32,441.45)=2741.68, p<.001; Semantic Decision: F(1.90,361.29)=521.94, p<.001; Spatial Context Probe: F(2.09,396.96)=424.98, p<.001; Semantic Probe: F(2.07,393.80)=117.88, p<.001; Greenhouse-Geisser correction applied). In follow-up contrasts using paired t-tests we compared the average Pearson r correlation of each phase to its relevant pathway, contrasted with the two non-pathway conjunctions. Correlations between the semantic probe and decision phases’ univariate activation and the semantic pathway were higher than non-pathway correlations and an equivalent pattern was seen for the spatial context pathway (Figure 4g; for exact t and p values associated with these comparisons see Supplementary Table S2), confirming that dissociable visual to DMN responses associated with semantic and spatial cognition are reliably present for individual participants.

Next, we examined if these pathways were reflected in the strength of white matter tracts connecting visual and DMN regions (seeds in Figure 4a). We examined structural connectivity in a subset of the HCP dataset (n = 164), asking if object probe visual regions showed a greater proportion of white matter tracts terminating in semantic DMN regions, and if scene probe visual regions showed stronger structural connectivity to spatial context DMN regions (Figure 4, panel h). A 2×2 repeated-measures ANOVA with visual regions as seeds and DMN regions as ROIs revealed a significant main effect of seed (F(1,163)=5.13, p=.025), ROI (F(1,163)=82.46, p<.001) and their interaction (F(1,163)=664.57, p<.001). Post-hoc comparisons confirmed stronger structural connectivity from the semantic probe visual regions to the semantic decision DMN regions; likewise, the spatial context probe visual regions showed stronger structural connectivity to the spatial context decision DMN regions (semantic probe visual: t(163)=8.25, p<.001; context probe visual: t(163)=478.66, p<.001). Repeating this analysis using the decision DMN regions as seeds and the probe visual regions as ROIs revealed a similar pattern (see Supplementary Analysis: “Replicating pathways’ structural connectivity from the DMN end” and Figure S6).

Finally, we examined how connectivity within the pathways changes depending on task demands in a Psychophysiological Interaction (PPI) analysis (see supplementary materials). We took the visual regions showing differential activation to object and scene probes as seeds (shown in Figures 2e and 4a), while the ROIs were regions sensitive to semantic and spatial context decisions within the DMN (shown in Figures 3e and 4a). The results, shown in Supplementary Figure S7, showed that the object seed was more connected to both semantic and spatial context DMN decision regions during the semantic task, while the scene probe regions were more connected to spatial context decision regions during the spatial context task than object probe regions.

#### Location of pathways in whole-brain gradients

Having found evidence for dissociable semantic and spatial context pathways, we analysed their location in a functional state space defined by the first two gradients of intrinsic connectivity (Margulies et al., 2016). The principal gradient relates to connectivity differences between unimodal and heteromodal cortex, while the second gradient captures connectivity differences between visual and auditory/somato-motor cortex. By locating the ends of the two visual-to-DMN pathways within gradient space, we can establish if DMN regions supporting semantic and spatial cognition are equally distant in connectivity from sensory-motor cortex: semantic cognition is arguably more abstract than spatial cognition and might be supported by DMN regions that are more isolated from sensory-motor systems on the principal gradient (Margulies et al., 2016; Smallwood et al., 2021). We can also establish if semantic and spatial DMN regions differ in the balance of connectivity to visual versus auditory-motor regions on the second gradient: heteromodal concepts are thought to be constructed from diverse sensory-motor features (Lambon Ralph et al., 2017), while spatial representations might draw more strongly on visual information (Epstein and Baker, 2019). We tested these predictions by locating individual unthresholded peak response coordinates for semantic and spatial context probes (within visual networks) and decisions (within the DMN) in gradient space (masked by Yeo et al.’s 7 network parcellation). We then asked if there are significant differences in the gradient locations of these tasks across participants.

The results (Figure 5, panel a) showed that there were no differences between the two tasks during the probe phase, while the decision phase was associated with task effects: DMN peaks for semantic decisions were more distant from sensory-motor cortex on the principal gradient, compared with spatial context decisions (t(1,26)=2.34, p=.027), consistent with the view that semantic cognition draws on more abstract and heteromodal representations in DMN. In addition, responses for the spatial context task were closer to the visual end of the second gradient, while responses for the semantic task were somewhat more balanced across visual and auditory-motor ends of this gradient (t(1,26)=3.31, p=.003)^2^. Since the scatterplots in Figure 5 do not distinguish whether these effects took place at the individual level (the data points are not linked across tasks), we plotted the same data comparing the gradient values for the peak responses in each of our tasks at the participant level in the Supplementary Materials (see Figure S8). This plot shows that in the majority of individual cases, the pattern of group-level results shown in Figure 5 held.

#### Cross-pathway integration of semantic and spatial cognition: Response to Same- versus Mixed-Category Buildings

Having identified dissociable semantic and spatial context pathways, and examined how these are differentially recruited across tasks, we investigated the integration of semantic and spatial context information across these processing streams. We compared responses in SCB and MCB trials, since semantic and spatial information are aligned when buildings contain items from a single semantic category, but not in mixed-category buildings. In these analyses, there were differences between conditions in the probe but not the decision-making phase (perhaps because many probes were presented without decisions, increasing statistical power).

First, we performed univariate contrasts of MCB and SCB trials (Figure 6, panels a-c). For scene probes in the spatial context task, the MCB > SCB contrast elicited a stronger response in dorsal lateral occipital cortex and retrosplenial cortex (Figure 6, panel a). In these circumstances, spatial context probes could only activate spatial and not semantic information. The SCB > MCB contrast activated an adjacent region of right angular gyrus (Figure 6, panel b). For semantic probes, the contrast of MCB > SCB identified greater engagement in distributed parietal, occipital and temporal regions, associated with the multiple-demand network (Figure 6, panel c). There were no clusters that showed a stronger response to same than mixed category building probes for the semantic task. There was also an interaction between task and condition, which was driven by the SCB > MCB effect in right angular gyrus in the spatial context task exceeding this effect in the semantic task.

To interpret these results, we conducted seed-to-ROI intrinsic connectivity analysis using independent data from Study 3, taking these clusters as seeds and the semantic and spatial context pathways masks (shown in Figure 4d and Figure 6, panel d) as ROIs. A two-way repeated measures ANOVA examined seed (spatial context SCB>MCB and MCB>SCB; semantic MCB>SCB) and ROI (semantic and spatial context pathways) as factors. The results can be seen in Figure 6, panel e). There were significant effects of seed (F(1.92,364.1)=211.48, p<.001), ROI (F(1,190)=182.67, p<.001) and an interaction (F(1.84,349.66)=723.412, p<.001). Post-hoc comparisons showed the context pathway was most connected to the spatial context MCB>SCB clusters, less connected to the spatial context SCB>MCB (when semantic information was also relevant to the response) and least connected to the semantic MCB>SCB regions (context MCB > context SCB: t(1,190)=34.91, p<.001; context SCB > semantic MCB: t(1,190)=5.18, p<.001). The opposite pattern of connectivity was found for the semantic pathway, which was most connected to the semantic MCB>SCB regions, less connected to the spatial context SCB>MCB and least connected to the spatial context MCB>SCB clusters (semantic MCB > context SCB: t(1,190)=16.93, p<.001; context SCB > context MCB: t(1,190)=10.59, p<.001). In this way, spatial context SCB>MCB regions, which reflected the engagement of semantic information in a spatial context task, showed an intermediate pattern of connectivity to both pathways.

A supplementary analysis of the task using multivariate approaches (see supplementary materials “Supplementary Analysis: Multivariate Response to Same- versus Mixed-Category Buildings”) found a similar pattern. We performed Representational Similarity Analysis (RSA) using a searchlight approach, which allowed us to detect regions sensitive to semantic and spatial context information. The results of this analysis in the MCB trials, where information could not be integrated, showed regions sensitive to category in the semantic task in bilateral ventral LOC, as well as regions sensitive to location in the spatial context task, dorsally in left LOC (see Supplementary Figure S9). A cross-task RSA of the SCB trials, where semantic and spatial information were aligned, allowing integration, identified a separate set of regions in right LOC, topographically situated between the two pathways, that captured spatial context information during the semantic task. An intrinsic connectivity analysis in data from Study 3 using the RSA regions as seeds and the pathways as ROIs showed that the regions sensitive to semantic and spatial context information during MCB trials were maximally connected to the semantic and spatial context pathways, respectively. On the other hand, the regions where spatial information could be decoded during the SCB trials of the semantic task showed an intermediate pattern of connectivity to both pathways (especially to the semantic pathway), suggesting a role in integrating information between them (see Figure S9).

## Discussion

Functional subdivisions of visual cortex and DMN sit at opposing ends of parallel processing streams supporting visually-mediated object-centric semantic and spatial cognition; moreover, regions with intermediate patterns of connectivity are implicated in the integration of these streams into coherent experience. Viewing object probes in a semantic task and location probes in a spatial context task activated different parts of visual cortex, functionally related to the passive viewing of objects and scenes. Semantic and spatial context decisions about these probes engaged FT and MT-DMN subsystems. Visual regions sensitive to object probes showed stronger intrinsic functional connectivity and structural connectivity to FT-DMN, while scene probe regions were more connected to MT-DMN. In a functional space defined by whole-brain connectivity patterns, the object-centric semantic pathway was more distant from unimodal regions on a unimodal-to-heteromodal connectivity gradient, and it had a more balanced influence of visual and auditory-motor systems, while the spatial context pathway was more visual. Finally, we found evidence that both heteromodal and visual regions integrate information about meaning and spatial context. When all the items in a building were drawn from a particular semantic category, there was greater recruitment of right angular gyrus; multivariate pattern analysis similarly found a cluster in lateral occipital cortex that represented spatial context information during the semantic task. When there was no opportunity to integrate object-centric semantic information with spatial context, regions that responded showed higher pathway-specific connectivity. In contrast, when integration was facilitated by the structure of the task, response regions had an intermediate pattern of connectivity.

Our study has important implications for the organisation of DMN into specialised subsystems, and for how these subsystems get their input from perceptual regions. Previous literature has robustly established distinct FT and MT subsystems (Andrews-Hanna et al., 2014a; Andrews-Hanna and Grilli, 2021a; Smallwood et al., 2021; Yeo et al., 2011); however, the way in which this architecture reflects differences in visual inputs remains contentious. One proposal is that different DMN subnetworks are differently engaged by tasks that are externally versus internally oriented. For example, (Chiou et al., 2020a) propose that there is a basic distinction between parts of the network that process semantic information accessed from words and images, and between DMN regions that sustain internally-focussed cognition. Other work has called into question whether semantic responses in FT-DMN are specific to external tasks: for example, (Zhang et al., 2022) found that lateral temporal regions changed their patterns of connectivity depending on the task, with more visual connectivity in externally-oriented tasks like reading, and more DMN connectivity in internally-orientated conceptual states like mind-wandering and autobiographical memory. This work suggests FT-DMN might support object-centric semantic cognition across internal and external modes of cognition.

Our findings also suggest that the distinction between these subsystems is not organised according to visual coupling; instead, DMN organisation arises from differential connectivity between distinct visual and DMN regions that gives rise to partially segregated pathways that process information about locations and meanings. Visual responses to scenes and objects reflect entry points to these processing pathways such that the key distinction between FT-DMN and MT-DMN relates to the type of information being processed, as opposed to how the information is accessed.

Our observed dissociation between semantic and spatial context pathways echoes a similar domain-specific organisation for working memory in prefrontal cortex (Levy and Goldman-Rakic, 2000; Romanski, 2004), in which there are dorsal and ventral streams associated with the maintenance of item location and identity respectively. This organising principle has been extended to long term memory more recently. (Deen and Freiwald, 2021) found a similar dissociation between places and people (instead of objects) in the DMN and other areas of association cortex. This was not tied to a specific input modality or task, indicative of parallel, domain-specific networks, at the top of the cortical hierarchy. Here, we extend this approach to consider whether functional divisions within DMN and visual cortex are connected, giving rise to pathways which are differentially situated in a connectivity state space defined by whole-brain dimensions of intrinsic connectivity, and we ask how these pathways might be integrated, and how they might be flexibly recruited according to task demands.

Although our research suggests a *domain*-specific view of brain organisation within visual-to-DMN pathways linked to object-centric semantic and spatial cognition, there may be different *processes* within meaning and spatial context tasks that drive these effects. The FT subsystem is thought to rely on the abstraction of information from sensory-motor inputs (Chiou et al., 2020a; Smallwood et al., 2021; Wang et al., 2020). The MT subsystem, on the other hand, uses a relational code that can capture spatial relations to successfully navigate complex environments (Eichenbaum, 2004; Eichenbaum and Cohen, 2014; Zeidman et al., 2015). One possibility is that, at the visual end of these pathways, spatial location is more dependent on peripheral vision, while object recognition is dependent in central fixation (Hasson et al., 2002; Levy et al., 2001); consequently, the distinct visual-to-DMN pathways we have recovered may reflect a basic property of how peripheral and central visual regions project to DMN. Our findings mirror and extend the results of Silson and colleagues (Silson et al., 2019; Steel et al., 2021), since we identify dissociable pathways between visual cortex and DMN; however, we extend this work to cover fully distributed networks that support object-centric semantic and spatial decision-making, and locate these pathways in a whole-brain gradient space relating to variation in patterns of intrinsic connectivity, as well as considering how these pathways can be integrated.

One question remains: how does the brain generate a coherent, seamlessly integrated experience of place and the identity of objects from these segregated, specialised streams of processing? The response we identified in right angular gyrus when object semantic and spatial context information was aligned is consistent with earlier studies implicating this brain region in the integration of information from multiple domains into a rich, meaningful context that can guide ongoing cognition (Lanzoni et al., 2020). One recent proposal suggests that neurons in this area represent high-dimensional inputs on a low-dimensional manifold encoding the relative position of items in physical space and abstract conceptual space (Summerfield et al., 2020). This region, which is maximally distant from sensory-motor cortex and equidistant from visual and motor cortex, might have the capacity to form representations that are not dominated by one type of input or code. We found a region of the right AG that was potentially important for integrating semantic and spatial context information. Previous research has established a key role of the AG in context integration (Bonnici et al., 2016; Branzi et al., 2020; Ramanan et al., 2017) and specifically, in guiding multimodal decisions and behaviour (Humphreys et al., 2021; Xu et al., 2017; Yazar et al., 2017). Although some recent proposals suggest a causal role of right AG in the early establishment of meaningful contexts, allowing semantic integration across modalities (Bocca et al., 2015; Muggleton et al., 2008; Olk et al., 2015; Petitet et al., 2015; Seghier, 2023), the majority of this research points to left, rather than right, AG as a key region for integration. We might have observed involvement of the right AG in our study since people were integrating semantic and spatial information, and visuospatial memory processes might be somewhat right lateralised (cf. Sormaz et al., 2017) and more strongly connected to right than left AG. We are not aware of a literature on right AG lesions impairing the integration of semantic and spatial information but, in the face of our findings, this might be a promising new direction. Patients with damage to right AG should be examined with specific tasks aimed at probing this type of integration. We also found evidence of information integration in occipital regions that were closer to the input regions of the visual to DMN pathways. These different levels of integration shared a common characteristic: in both cases, the region implicated in integration was spatially interposed between the pathways, consistent with the view that topography is highly relevant to information integration since adjacent brain regions tend to share a high degree of functional connectivity and represent similar information.

While we might assume that common visual-to-DMN pathways support memory access from vision (as in this study), and subserve the generation of visual features when imagining objects versus scenes, this hypothesis awaits empirical investigation. Moreover, our pathways are vision-specific, and it remains unclear if there are analogous pathways from auditory or somatomotor cortex to DMN. The generality of these pathways must be confirmed across tasks, since spatial representations are likely to interact with other representational codes, including emotion and social information – the interdigitated pathways highlighted in these circumstances (Braga and Buckner, 2017; DiNicola et al., 2020) might show anatomical differences or be broadly the same as the pathways uncovered here.

Likewise, further research should be carried out on memory-visual interactions for alternative domains. Our study focused on spatial location and semantic object processing and therefore cannot address how other categories of stimuli, such as faces, are processed by the visual-to-memory pathways that we have identified. Previous work has suggested some overlap in the neurobiological mechanisms for semantic and social processing (Andrews-Hanna et al., 2014b; Andrews-Hanna and Grilli, 2021b; Chiou et al., 2020b), suggesting that the FT-DMN pathway may be highlighted when contrasting both social faces and semantic objects with spatial scenes. On the other hand, some researchers have argued for a ‘third pathway’ for aspects of social visual cognition (Pitcher, 2023; Pitcher and Ungerleider, 2021). Future studies that probe other categories will be able to confirm the generality (or specificity) of the pathways we described.

One important caveat is that we have not investigated the spatiotemporal dynamics of neural propagation along the pathways we identified between visual cortex and DMN. The dissociations we found in task responses, intrinsic functional connectivity and white matter connections all support the view that there are at least two distinct routes between visual and heteromodal DMN regions, yet this does not necessarily imply that there is a continuous sequence of cortical areas that extend from visual cortex to DMN – and given our findings of structural connectivity differences that relate to the functional subdivisions we observe, this is unlikely to be the sole mechanism underpinning our findings. It would be interesting in future work to characterise the spatiotemporal dynamics of neural propagation along visual-DMN pathways using methods optimised for studying the dynamics of information transmission, like Granger causality or travelling wave analysis.

Moreover, many questions remain about information integration across the semantic and spatial domains; does spatial juxtaposition promote the emergence of an integrated code, and are neural representations that emerge at the intersection of these pathways more than the sum of their parts? Although further research is needed, the current study highlights how subdivisions within visual and DMN networks are related to types of information, giving rise to distinct processing streams that capture different unimodal to heteromodal transformations relevant to object-centric semantic and spatial context processing, and shows how these pathways might interact at multiple levels of the cortical hierarchy to produce coherent cognition.

## Methods

### Study 1. Task-based fMRI

#### Study 1: Participants

Thirty native English speakers (mean age = 22.6 ± 2.7 years, age-range 18-34 years, 8 males) with normal or corrected-to-normal vision and no history of language disorders participated in this study. Ethical approval was obtained from the Research Ethics Committees of the Department of Psychology and York Neuroimaging Centre, University of York. Written informed consent was obtained from all subjects prior to testing.

#### Study 1: Materials

The learning phase employed videos showing a walk-through for twelve different buildings (one per video), shot from a first-person perspective. The videos and buildings were created using an interior design program (Sweet Home 3D). Each building consisted of two rooms: a bedroom and a living room/office, with an ajar door connecting the two rooms. The order of the rooms (1st and 2nd) was counterbalanced across participants. Each room was distinctive, with different wallpaper/wall colour and furniture arrangements. The building contexts created by these rooms were arbitrary, containing furniture that did not reflect usual room distributions (i.e., a kitchen next to a dining room), to avoid engaging further conceptual knowledge about frequently-encountered spatial contexts in the real world. Within each room, there were three framed images of objects and animals, towards the start, middle, and end of each video (see top panel of Supplementary Figure S1), each at the same distance from its neighbour (across the two rooms); seventy-two images were presented overall. The images represented single items from 6 semantic categories: musical instruments, gardening tools, sports equipment, mammals, fish and birds (12 pictures from each category). Half of the buildings contained images from the same semantic category (‘Same Category Building’, SCB), and the other half contained images from different semantic categories (‘Mixed Category Building’, MCB). The presentation of items within each room in Mixed Category Buildings was controlled such that: 1) no item from the same category was presented at the same location twice, and 2) no three categories were grouped together more than once.

A full list of pictures of the object and location stimuli employed in this task, as well as the videos watched by the participants can be consulted in the OSF collection associated with this project under the components OSF>Tasks>Training.

#### Study 1: Design and Procedure

*Training Task:* Subjects participated in a training session the day before the MRI scan, where they watched the walkthrough videos, each lasting 49 s (Figure 9 and top panel of Figure S1). Participants watched each video at least 6 times in 3 rounds (twice per round). Each round consisted of 4 mini-blocks of videos containing 3 videos. After each mini-block, participants were given a test in Psychopy3: they were asked to choose the room that each item was presented in, responding via button press. Items were pictured at the top of the screen, with the correct room and another room below (Supplementary Figure S1, left half of bottom panel). Screenshots were taken of the location of each framed image and the rooms themselves (from the entrance way), with the images and their frames removed (see bottom panel of Supplementary Figure S1). They had 5 s to respond, after which the correct room was presented as feedback for a further 5 s. Following each round, there was a matching task, which reinforced participants’ memory of which rooms belonged together. Two rooms from the same building were presented, with two items from that building (one from each room) below. Participants were instructed to drag the objects into the correct room of the building (Supplementary Figure S1, right half of bottom panel). Feedback showed the correct object in each room. Finally, to establish how well the item-location pairs were learned, participants were given a final test on all the rooms and items: this followed the structure of the mini-block tests, except that materials from the entire session were included. If accuracy was below 80%, participants watched the videos again until this threshold was reached. In total participants spent approximately 2 hours on the training. The amount of training required was established in pilot testing with 9 participants who did not take part in the main study. This also confirmed the items were easily nameable.

**Figure 9.**
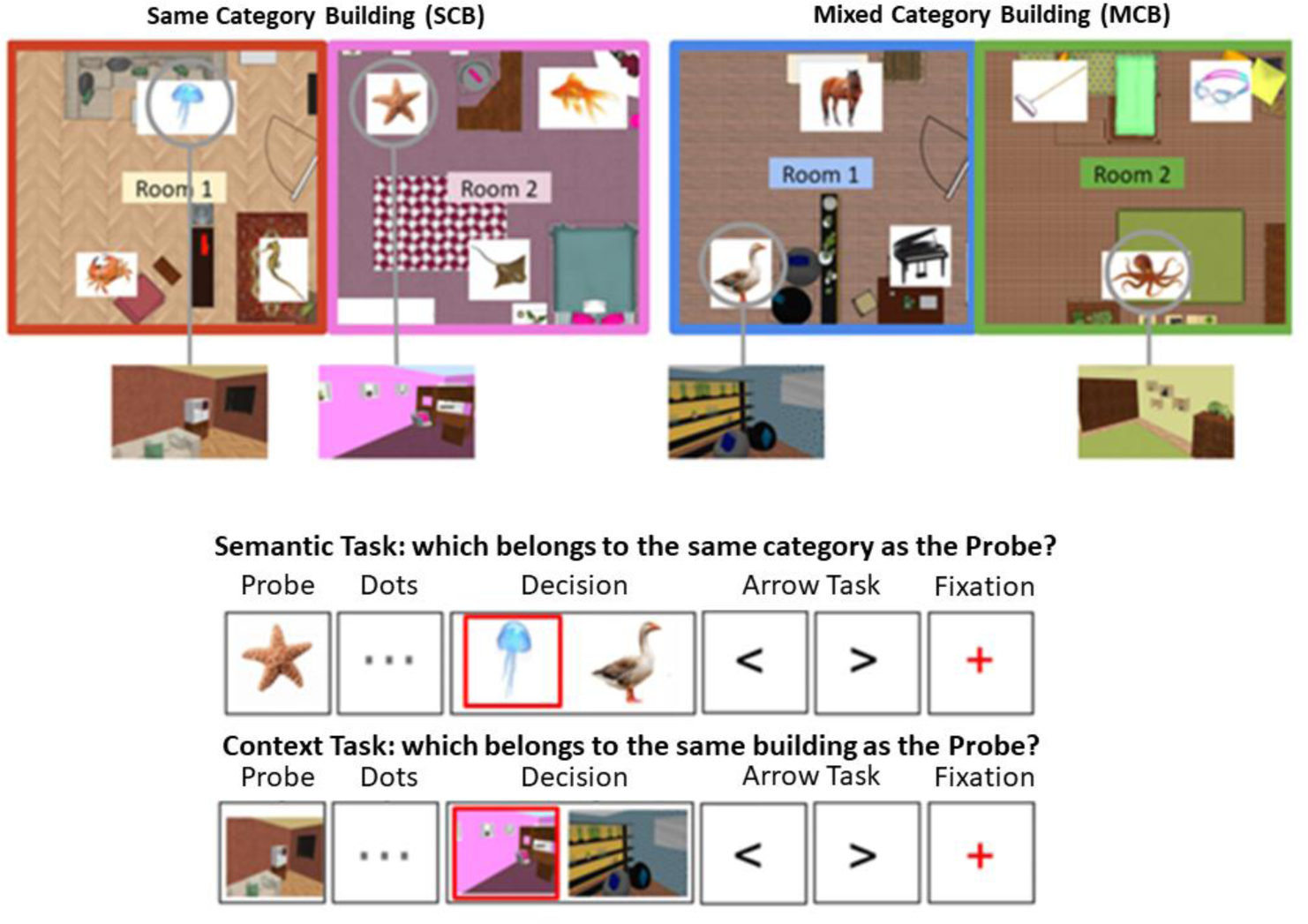
Top panel: the layout of two buildings is shown, one of them contains semantically related items (SCB), the other contains unrelated items (MCB). These items and locations are shown in the example trials below. Bottom panel: trial procedure for Semantic and Spatial Context decisions. The phases of a trial are shown (Probe, Dots, Decision, Arrow task, Fixation), and the red square indicates the correct response (not shown to participants). Participants were required to press ‘left’ or ‘right’ buttons in the Decision phase. No-decision trials omitted the dots and decision phases.

*fMRI Task:* On the day of the scan, participants repeated the final test from the training day to establish how well they retained the information. They then watched all 12 videos once again, in a counterbalanced order, and performed the test phase a second time. The mean accuracy on the first day was 95.1%, (SD=5.7%) while on the second day it was 97.1% (SD=3.8%)^3^.

Inside the scanner, participants performed a semantic and spatial context memory task, using a slow event-related design (Figure 9). The semantic task involved judgements about the semantic category of objects and animals (using the same images that has been presented in the buildings), while the spatial context task involved matching rooms that belonged to the same building. Both tasks consisted of ‘no-decision’ and ‘decision’ trials. No-decision trials were optimised for representational similarity analysis and included an image of the probe item (for 2s) followed by an “arrow task” in which participants pressed ‘left’ or ‘right’ to match the direction of a series of chevrons (‘<’ ‘>’) presented on the screen (for 3s), ending with a red fixation cross (1s). The probe in the semantic task was an object or animal from the training; for the spatial context task, it was a screenshot of an item’s location within a room (excluding the item itself). In decision trials, the same types of probes were presented but they were followed by three central dots (for 4s) indicating a decision would be made (Figure 9). In the decision phase, a target image (from the same category or building as the probe) was presented together with a distractor image, creating a 2-alternative forced-choice judgment. In the spatial context task, the distractor was a room from a different building, while in the semantic task, the distractor was an item from a different category. On SCB (same category building) trials, semantic and spatial information were aligned in the sense that the target and probe were from the same category and the same building. On MCB (mixed category building) trials, semantic and spatial information did not converge: semantic targets were in a different building from the probe, while spatial context targets were from a different semantic category from the probe (see Figure 9 for the structure of semantic and spatial context trials). Decisions were required within 4 s, and then the task progressed to the next trial. Participants made their responses using a button box, pressing with their right index finger to indicate whether the target image was on the left or right side of the screen. Participants were encouraged to respond as quickly and accurately as possible. After the decision was made, participants carried out the arrow task again (for 6.5s minus their response time) followed by a red fixation cross (1s) indicating the end of the trial. The arrow task served as a non-memory baseline and was included to increase separation of the BOLD signal between trials.

During the fMRI scan, there were 4 runs of the spatial context task and 4 runs of the semantic task. Each run contained 36 trials and lasted approximately 6 minutes. All 72 objects were presented as stimuli across blocks 1 and 2, and across blocks 3 and 4. Each run included 18 decision and 18 no decision trials. The decision trials in each run were further subdivided into 9 SCB and 9 MCB decision trials. The decision trials in run 1 and run 2 became the no-decision trials in runs 3 and 4, and vice versa. Spatial context runs preceded semantic runs. The order of trials within each run was counterbalanced between participants. Prior to scanning, participants were given formal instructions for the tasks and shown how to use the response box.

#### Study 1: Task-based fMRI

*MRI Data Acquisition:* Whole brain structural and functional MRI data were acquired using a 3T Siemens MRI scanner utilising a 64-channel head coil, tuned to 123 MHz at York Neuroimaging Centre, University of York. A Localiser scan and 8 whole-brain functional runs (4 of the semantic task, 4 of the spatial context task) were acquired using a multi-band multi-echo (MBME) EPI sequence, each approximately 6 minutes long (TR = 1.5 s; TEs = 12, 24.83, 37.66 ms; 48 interleaved slices per volume with slice thickness of 3 mm (no slice gap); FoV = 24 cm (resolution matrix = 3×3×3; 80×80); 75° flip angle; 705 volumes per run (235 TRs with each TR collecting 3 volumes); 7/8 partial Fourier encoding and GRAPPA (acceleration factor = 3, 36 ref. lines; multi-band acceleration factor = 2). Structural T1-weighted images were acquired using an MPRAGE sequence (TR = 2.3 s, TE = 2.26 s; voxel size = 1×1×1 isotropic; matrix size = 256 x 256, 176 slices; flip angle = 8°; FoV= 256 mm; ascending slice acquisition ordering).

*Multi-echo Data Pre-processing:* This study used a multiband multi-echo (MBME) scanning sequence to optimise signal from medial temporal regions (e.g., ATL, MTL) while also maintaining optimal signal across the whole brain (Halai et al., 2014). We used TE Dependent ANAlysis (TEDANA, version 0.0.10, https://doi.org/10.5281/zenodo.4725985, https://tedana.readthedocs.io/) to combine the images (Kundu et al., 2013; Posse et al., 1999). Before images were combined, some pre-processing was performed. FSL_anat (https://fsl.fmrib.ox.ac.uk/fsl/fslwiki/fsl_anat) was used to process the anatomical images, including re-orientation to standard (MNI) space (fslreorient2std), automatic cropping (robustfov), bias-field correction (RF/B1 – inhomogeneity-correction, using FAST), linear and non-linear registration to standard-space (using FLIRT and FNIRT), brain extraction (using FNIRT, BET), tissue-type and subcortical structure segmentation (using FAST). The multi-echo data were pre-processed using AFNI (https://afni.nimh.nih.gov/), including de-spiking (3dDespike), slice timing correction (3dTshift; heptic interpolation), and motion correction of all echoes aligned to the first echo (with a cubic interpolation; 3dvolreg was applied to the first echo to realign all images to the first volume; these transformation parameters were then applied to echoes 2 and 3). The pre-processing script is available at OSF (https://osf.io/sh79m/).

*Task-based fMRI Data Analysis:* Further pre-processing of the functional and structural data was carried out using FSL version 6.0 (Jenkinson et al., 2002; Smith et al., 2004; Woolrich et al., 2009). Functional data were pre-processed using FSL’s FMRI Expert Analysis Tool (FEAT). The TEDANA outputs (denoised optimally combined timeseries) registered to the participants’ native space were submitted to FSL’s FEAT. The first volume of each functional scan was deleted to negate T1 saturation effects. Pre-processing included high-pass temporal filtering (Gaussian-weighted least-squares straight line fitting, with sigma = 50s), linear co-registration to the corresponding T1-weighted image followed by linear co-registration to MNI152 2mm standard space (Jenkinson and Smith, 2001), which was then further refined using FSL’s FNIRT nonlinear registration (J L R Andersson et al., 2007; Jesper L. R. Andersson et al., 2007) with 10mm warp resolution, spatial smoothing using a Gaussian kernel with full-width-half-maximum (FWHM) of 5 mm, and grand-mean intensity normalisation of the entire 4D dataset by a single multiplicative factor.

*Task GLM:* Second and group-level analyses were also conducted using FSL’s FEAT version 6. Pre-processed time series data were modelled using a general linear model in FSL, using FILM correcting for local autocorrelation (Woolrich et al., 2001). We used an event-related design. Our aim was twofold: (1) to characterise differential activation between the semantic and spatial context tasks at each phase of the trials, and (2) to document any potential differences of activation in response to MCB and SCB trials in probe and decision phases, in each task. To this end, the following 8 EVs were entered into a general linear model, convolved with a double-gamma haemodynamic response function: the probe, dots and decision phases (only correct responses) were modelled for both MCB and SCB trials (3×2 EVs). Correct decisions made during the arrow task were modelled in a separate EV to use as an explicit baseline. Incorrect and omitted responses in the decision phase, as well as errors made during the arrow task, were combined into a regressor of no interest. The fixation crosses between trials were not explicitly modelled. Probe and dots phases were modelled as fixed-duration epochs, while semantic, spatial context and arrow decisions were modelled using a variable epoch approach, based on each participant’s reaction time on that trial. At the first level, the semantic and spatial context tasks were entered into separate models for each run performed by all participants. We then combined all valid runs for each participant into a participant level analysis, again separately for each task, at the second level (see ‘Data Exclusions’ below for details).

At the group level, we performed two separate univariate analyses. First, we compared activation for the two tasks, contrasting semantic and spatial context models. Inputs for this analysis were lower-level contrasts of the probe phase of each task against the implicit baseline, and the decision phase of each task contrasted against the explicit baseline of arrow decisions. In our second analysis, we used the same lower-level contrasts but examined the semantic and spatial context tasks separately, examining within-task differences between MCB and SCB trials in the probe and decision phases. This also allowed us to explore interactions between MCB/SCB trials and task. We did not include any motion parameters in the model as the data submitted to these first level analyses had already been denoised as part of the TEDANA pipeline (Kundu et al., 2012). At the group-level, analyses were carried out using FMRIB’s Local Analysis of Mixed Effects (FLAME1) stage 1 with automatic outlier detection (Beckmann et al., 2003; Woolrich, 2008; Woolrich et al., 2004), using a (corrected) cluster significance threshold of p = 0.05, with a z-statistic threshold of 2.6 (Eklund et al., 2016) to define contiguous clusters.

*Data Exclusions:* We excluded three participants: one due to excessive motion (mean framewise displacement > 0.3mm) in more than 50% of functional runs, another due to misunderstanding the task (0% accuracy in MCB decisions in 3 out of 4 runs of the semantic task), and one due to low SCB accuracy in 3 out of 4 runs of the spatial context task, with less than 50% of usable data. We also excluded any individual runs where the decision accuracy was equal or below chance level (50%) in the SCB condition^4^. This led to the removal of four runs across three participants in the semantic task, and twelve runs across eight participants in the spatial context task. Two runs were removed due to data loss (a corrupted EV file and data transfer failure from the MRI scanner). 92.5% of the runs acquired were included in the analysis.

#### Study 1: Psychophysiological Interaction Analysis

In order to test for distinct semantic and spatial memory pathways that connect visual regions to distinct subnetworks of the DMN, we conducted a psychophysiological interaction (PPI) supplementary analysis. In short, we created semantic and spatial context seeds from the visual regions activated to object and scene probes, and examined their connectivity to the DMN regions activated during the decision phase of the tasks, using two separate models (one for each seed) which examined the main effect of the task. We describe the methods in detail in the relevant section of the Supplementary Materials (Supplementary Analysis: Effects of task demands on pathway connectivity).

#### Study 1: Representational Similarity Analysis

Since distinct but adjacent regions were associated with semantic and spatial context decisions, we asked what they represented during probe presentation using Representational Similarity Analysis (RSA). We constructed semantic similarity matrices where trials that shared a specific category (e.g., birds) were assigned the strongest value, and spatial context similarity matrices where pairs of trials belonging to the same room were assigned the strongest value. After single-trial estimation using a Least Square-Single approach, we carried out second-order RSA using a searchlight approach to compare semantic and spatial context similarity matrices with neural similarity matrices. This allowed us to identify voxels that were sensitive to semantic and spatial relationships between probes in each of our tasks. We also performed cross-task similarity analysis, correlating semantic similarity to the neural similarity matrix from the spatial context task (and vice versa), to identify regions sensitive to semantic and spatial context information across tasks. We report the methods in detail in the section for this analysis in the Supplementary Materials (Supplementary Analysis: Multivariate Response to Same- versus Mixed-Category Buildings).

### Study 2. Passive Viewing of Objects and Scenes

We examined passive viewing of objects and scenes in a sample of fifty-two healthy volunteers, providing independent regions-of-interest for the analyses of Studies 1 and 3.

#### Study 2: Participants

Fifty-two participants with normal, or corrected-to-normal, vision gave informed consent. The study was approved by the Research Ethics Committee at York Neuroimaging Centre.

#### Study 2: Stimuli

Dynamic stimuli were 3-second movie clips of faces, bodies, scenes, objects and scrambled objects (see Supplementary Figure S2) designed to localize category-selective visual areas (Pitcher et al., 2011). Only the scenes and object stimuli were used in the present study. There were sixty movie clips for each category in which distinct exemplars appeared multiple times. Fifteen different locations were used for the scene stimuli which were mostly pastoral scenes shot from a car window while driving slowly through leafy suburbs, along with films flying through canyons or walking through tunnels that were included for variety. Fifteen different moving objects were selected that minimized any suggestion of animacy of the object itself or of a hidden actor pushing the object (these included mobiles, windup toys, toy planes and tractors, balls rolling down sloped inclines). Within each block, stimuli were randomly selected from within the entire set for that stimulus category (faces, bodies, scenes, objects, scrambled objects).

#### Study 2: Procedure and Data Acquisition

Functional data were acquired over 6 block-design functional runs lasting 234 seconds each. Each functional run contained three 18-second rest blocks, at the beginning, middle, and end of the run, during which a series of six uniform color fields were presented for three seconds. Participants were instructed to watch the movies but were not asked to perform any overt task.

Imaging data were acquired using a 3T Siemens Magnetom Prisma MRI scanner (Siemens Healthcare, Erlangen, Germany) at the University of York. Functional images were acquired with a twenty-channel phased array head coil and a gradient-echo EPI sequence (38 interleaved slices, repetition time (TR) = 3 sec, echo time (TE) = 30ms, flip angle = 90%; voxel size 3mm isotropic; matrix size = 128 x 128) providing whole brain coverage. Slices were aligned with the anterior to posterior commissure line. Structural images were acquired using the same head coil and a high-resolution T-1 weighted 3D fast spoilt gradient (SPGR) sequence (176 interleaved slices, repetition time (TR) = 7.8 sec, echo time (TE) = 3ms, flip angle = 20 degrees; voxel size 1mm isotropic; matrix size = 256 x 256).

#### Study 2: Imaging Analysis

Functional MRI data were analyzed using AFNI (http://afni.nimh.nih.gov/afni). Images were slice-time corrected and realigned to the third volume of the first functional run and to the corresponding anatomical scan. All data were motion corrected and any TRs in which a participant moved more than 0.3mm in relation to the previous TR were discarded from further analysis. The volume-registered data were spatially smoothed with a 4-mm full-width-half-maximum Gaussian kernel. Signal intensity was normalized to the mean signal value within each run and multiplied by 100 so that the data represented percent signal change from the mean signal value before analysis.

Data from all runs were entered into a general linear model (GLM) by convolving the standard hemodynamic response function with the regressors of interest (faces, bodies, scenes, objects, and scrambled objects) for dynamic and static functional runs. Regressors of no interest (e.g., 6 head movement parameters obtained during volume registration and AFNI’s baseline estimates) were also included in the GLM. Data from all fifty-two participants were entered in a group whole brain analysis. Group whole brain contrasts were generated to quantify the neural responses across the experimental conditions. Scene-selective areas were defined using a contrast of dynamic scenes greater than dynamic objects, and object-selective areas were defined using a contrast of dynamic objects greater than scrambled objects, following convention (Epstein and Kanwisher, 1998; Malach et al., 1995). Activation maps were calculated using a t-statistical threshold of *p* = 0.001 and a cluster correction of 50 contiguous voxels (as these thresholds have been successfully used in other studies to characterise activation in the visual perception literature, e.g., (Nikel et al., 2022; Zimmermann et al., 2018). The whole-brain results are presented in Supplementary Figure S3.

### Study 3. Analysis of intrinsic functional connectivity using resting-state fMRI

The results of Study 1 suggested separate visual-DMN pathways recruited by semantic and spatial context tasks. To provide converging evidence for this ‘dual pathway’ architecture, we examined the intrinsic connectivity of sites identified in the univariate and RSA analyses in a separate sample.

#### Study 3: Participants

One hundred and ninety-one student volunteers (mean age=20.1 ± 2.25 years, range 18 – 31; 123 females) with normal or corrected-to-normal vision and no history of neurological disorders participated in this study. Written informed consent was obtained from all subjects prior to the resting-state scan. The study was approved by the ethics committees of the Department of Psychology and York Neuroimaging Centre, University of York. This data has been used in previous studies to examine the neural basis of memory and mind-wandering, including region-of-interest based connectivity analysis and cortical thickness investigations (Evans et al., 2020; Gonzalez Alam et al., 2018, 2022, 2019, 2021; Karapanagiotidis et al., 2017; Poerio et al., 2017; Turnbull et al., 2018; Vatansever et al., 2017; Wang et al., 2018).

#### Study 3: Pre-processing

Pre-processing and statistical analyses of resting-state data were performed using the CONN functional connectivity toolbox V.20a (http://www.nitrc.org/projects/conn; (Whitfield-Gabrieli and Nieto-Castanon, 2012) implemented through SPM (Version 12.0) and MATLAB (Version 19a). For pre-processing, functional volumes were slice-time (bottom-up, interleaved) and motion-corrected, skull-stripped and co-registered to the high-resolution structural image, spatially normalized to the Montreal Neurological Institute (MNI) space using the unified-segmentation algorithm, smoothed with a 6 mm FWHM Gaussian kernel, and band-passed filtered (.008 - .09 Hz) to reduce low-frequency drift and noise effects. A pre-processing pipeline of nuisance regression included motion (twelve parameters: the six translation and rotation parameters and their temporal derivatives), scrubbing (outlier volumes were identified through the composite artifact detection algorithm ART in CONN with conservative settings, including scan-by-scan change in global signal z-value threshold = 3; subject motion threshold = 5 mm; differential motion and composite motion exceeding 95% percentile in the normative sample) and CompCor components (the first five) attributable to the signal from white matter and CSF (Behzadi et al., 2007), as well as a linear detrending term, eliminating the need for global signal normalization (Chai et al., 2012; Murphy et al., 2009).

#### Seed Selection and Analysis

Intrinsic connectivity seeds were binarised masks derived from: (1) significant univariate clusters; and (2) significant effects identified in representational similarity analysis. For semantic and spatial probe effects, which characterised effects of visual perception, we created ROIs within Yeo et al.’s (2011) visual central and peripheral networks combined. For semantic and spatial context decisions, we identified regions within Yeo et al.’s (2011) combined DMN subnetworks. We also examined the intrinsic connectivity of regions activated by SCB versus MCB probes in Study 1. For representational similarity analyses, all voxels that survived thresholding at p<.05 in the MCB conditions for the semantic and context task, as well as the cross-task analyses were binarised and used as seeds. We excluded all non-grey matter voxels that fell within these masks.

#### Spatial Maps and Seed-to-ROI Analysis

We performed seed-to-voxel analyses convolved with a canonical haemodynamic response function for each of these seeds. At the group-level, analyses were carried out using CONN with cluster correction at p < .05, and a threshold of p-FDR = .001 (two-tailed) to define contiguous clusters. Seed to ROI connectivity was extracted for each participant and seed using REX implemented in CONN (Whitfield-Gabrieli and Nieto-Castanon, 2012), with percentage signal change as units. These values were then entered into a series of repeated-measures ANOVAs.

#### Cognitive Decoding

Connectivity maps were uploaded to Neurovault (Gorgolewski et al., 2015) https://neurovault.org/collections/13821/) and decoded using Neurosynth (Yarkoni et al., 2011). Neurosynth is an automated meta-analysis tool that uses text-mining approaches to extract terms from neuroimaging articles that typically co-occur with specific peak coordinates of activation. It can be used to generate a set of terms frequently associated with a spatial map. The results of cognitive decoding were rendered as word clouds using in-house scripts implemented in R. We excluded terms referring to neuroanatomy (e.g., “inferior” or “sulcus”), as well as the second occurrence of repeated terms (e.g., “semantic” and “semantics”). The size of each word in the word cloud relates to the frequency of that term across studies.

### Structural connectivity analysis

To provide converging evidence for parallel visual-to-DMN pathways, we performed tractography analysis using DTI data from an independent sample derived from the Human Connectome Project (HCP).

#### DTI pre-processing

We used data from a subgroup of 164 HCP participants who underwent diffusion-weighted imaging at 3 Tesla (Uǧurbil et al., 2013)http://www.humanconnectome.org/study/hcp-young-adult/). The imaging parameters were previously described in Uǧurbil et al. 2013, and involved acquiring 111 near-axial slices with an acceleration factor of 32, an isotropic resolution of 1.25 mm3, and coverage of the entire head. The diffusion-weighted images were obtained using 90 uniformly distributed gradients in multiple Q-space shells (Caruyer et al., 2013), and this process was repeated three times with different b-values and phase-encoding directions. We used a pre-processed version of this dataset, previously described (Karolis et al., 2019; Thiebaut de Schotten et al., 2020; Vu et al., 2015), that included steps to correct for susceptibility-induced off-resonance field, motion, and geometrical distortion.

We used StarTrack software (https://www.mr-startrack.com) to perform whole-brain deterministic tractography in the native DWI space. We applied an algorithm for spherical deconvolutions (damped Richardson-Lucy), with a fixed fibre response corresponding to a shape factor of α = 1.5 × 10–3 mm2.s−1 and a geometric damping parameter of 8. We ran 200 algorithm iterations. The absolute threshold was set at three times the spherical fibre orientation distribution (FOD) of a grey matter isotropic voxel, and the relative threshold was set at 8% of the maximum amplitude of the FOD (Thiebaut de Schotten et al., 2011). To perform the whole-brain streamline tractography, we used a modified Euler algorithm (Dell’Acqua et al., 2013) with an angle threshold of 45°, a step size of 0.625 mm, and a minimum streamline length of 15 mm.

To standardize the structural connectome data, we followed these steps: first, we converted the whole-brain streamline tractography into streamline density volumes, with the intensity corresponding to the number of streamlines crossing each voxel. Second, we generated a study-specific template of streamline density volumes using the Greedy symmetric diffeomorphic normalization pipeline provided by ANTs. This average template was created for all subjects. Third, we co-registered the template with a standard 1mm MNI152 template using the FLIRT tool in FSL to produce a streamline density template in the MNI152 space. Finally, we registered individual streamline density volumes to the template and applied the same transformation to the individual whole-brain streamline tractography using ANTs GreedySyn and the Trackmath tool in the Tract Querier software package (Wassermann et al., 2016). This produced whole-brain streamline tractography in the standard MNI152 space.

#### Tract extractions and ROI analysis

Our starting point for extracting semantic and spatial context pathway tracts was each participant’s whole-brain streamline tractography in MNI (1mm) space. We used the same univariate regions described in Section 2.3.3 as seeds (i.e., the seeds in the intrinsic connectivity analysis): these consisted of regions that were activated during the probe phase of each task, masked by Yeo’s visual networks, and regions that were activated during the decision phase of each task, masked by Yeo’s DMN networks. For each of our seeds, we used Trackvis (Wang and Benner, 2007) to extract all streamlines emerging from these regions as a volume, yielding one streamline group per seed per participant. Then, for each probe visual seed, we calculated what percentage of streamlines touched one decision DMN ROI or the other (activated by semantic and spatial context decisions; percentages adding to 100%); likewise, for each decision DMN seed, we calculated what percentage of streamlines touched either visual probe ROI (activated by object and scene probes; again adding to 100%). This allowed us to examine if the object probe regions were more connected to the semantic decision DMN regions, and if the scene probe regions were more connected to the spatial context DMN regions, in line with dual pathways.

### Situating the pathways in whole-brain gradients

We examined the position of the semantic and context pathways in a functional connectivity space defined by the first two dimensions of whole-brain intrinsic connectivity patterns, frequently referred to as “gradients”. The first dimension of this space relates to the distinction between the connectivity patterns of unimodal and heteromodal cortical regions, while the second dimension captures the separation of visual and auditory/somatomotor regions (Margulies et al., 2016). This analysis can reveal whether the semantic pathway shows more of a balance between visual and somatosensory/auditory modalities than the spatial context pathway, in line with view that concepts are heteromodal, abstracted from multiple sensory-motor features (Lambon Ralph et al., 2017). The analysis can also show whether the spatial context pathway is anchored in more visual portions of this functional space, in line with this modality’s importance for scene processing (Epstein and Baker, 2019).

First, we examined the univariate BOLD activation for each participant during the probe and decision phases of each task. The decision phase was contrasted with the arrow task baseline to control for low level motor responses. Next, we identified the MNI voxel location of the peak response for each participant: for activation during the decision phase, this was done within a mask of the DMN from the Yeo et al. (2011) 7-network parcellation, while for probe responses, we performed this analysis within the visual network of the same parcellation. We then fitted a sphere with a 5mm radius around this peak and used it as a ROI to extract the mean value in Margulies et al.’s (2016) maps for the two dimensions or gradients described above. The results were entered into a repeated measures 2×2 ANOVA with task and gradient as factors to establish whether the semantic and spatial context pathways differed in their location in this functional space.

### ROI-based ANOVA analyses

ROI-based analyses of activation and intrinsic connectivity in Studies 1 and 3 were performed using FSL’s “Featquery” tool for Study 1 and REX for Study 3, which we used to extract the percentage signal change within unweighted, binarised masks. The ANOVAs were carried out using IBM SPSS Statistics version 27. The results of post-hoc tests to interpret significant interactions were corrected for multiple comparisons using the Holm-Bonferroni method (Aickin and Gensler, 1996). All the p values reported in the Results section are Holm-Bonferroni adjusted p values.

### Visualisations of neural results

Brain maps were produced in BrainNet (Xia et al., 2013) using the extremum voxel algorithm, with the exception of slices depicted in Figures 2-4, which were produced in FSL Eyes (Figures 2 and 3) and MRIcroGL (Figure 4). Maps are provided in the following Neurovault collection: https://neurovault.org/collections/13821/.

## Supporting information

Revised Supplementary Materials

Response to Reviewers' Comments

## Acknowledgements

This project was funded by the European Research Council (ERC) under the European Union’s Horizon 2020 research and innovation programme (Project ID: 771863 – FLEXSEM to EJ; Project ID: 818521 - DISCONNECTOME to MTS; Project ID: 866533 to DSM). Additionally, this work was conducted in the framework of the University of Bordeaux’s IHU ‘Precision & Global Vascular Brain Health Institute - VBHI’, IdEx ‘Investments for the Future’ program RRI “IMPACT”, which received financial support from the France 2030 program.

## Data Availability

The scripts used in the presentation of the task, the analysis of the neuroimaging data and the visualisation of the results reported here can be consulted in the OSF collection associated with this paper (https://osf.io/sh79m/). We do not have sufficient consent for the public release of individual-level data; researchers wanting access to these data should contact the Research Ethics Committee of the York Neuroimaging Centre. Data will be released when this is possible under the terms of the UK and EU General Data Protection Regulations. Group-level brain maps used to produce the figures are available from the following Neurovault collection: https://neurovault.org/collections/13821/.

## Footnotes

1 The Yeo et al. parcellation labels these subnetworks of the DMN as Core DMN = DMN-A, FT-DMN = DMN-B, MT-DMN = DMN-C.

2 An ANOVA including task and condition (MCB versus SCB) replicated these task effects and found no effects of condition on the position of peak responses in gradient space.

3 Two participants were excluded from the mean accuracy calculation for day 2 due to data loss.

4 This threshold was not applied to the MCB condition, which was expected to elicit interference between semantic and spatial context information.

