## Supplementary material for "Visual to default network pathways: A double dissociation between semantic and spatial cognition": Revised Supplementary Materials

Study 1. Training and Test Materials

In the training session, participants navigated virtual environments populated with objects, with the aim of learning these objects' location in the environments. During this training, their memory was tested in a two-alternative forced-choice test and a matching task. Figure S1 shows some example screenshots from videos used by participants to learn the environments, and the tests used during the training session to probe participants' memory.

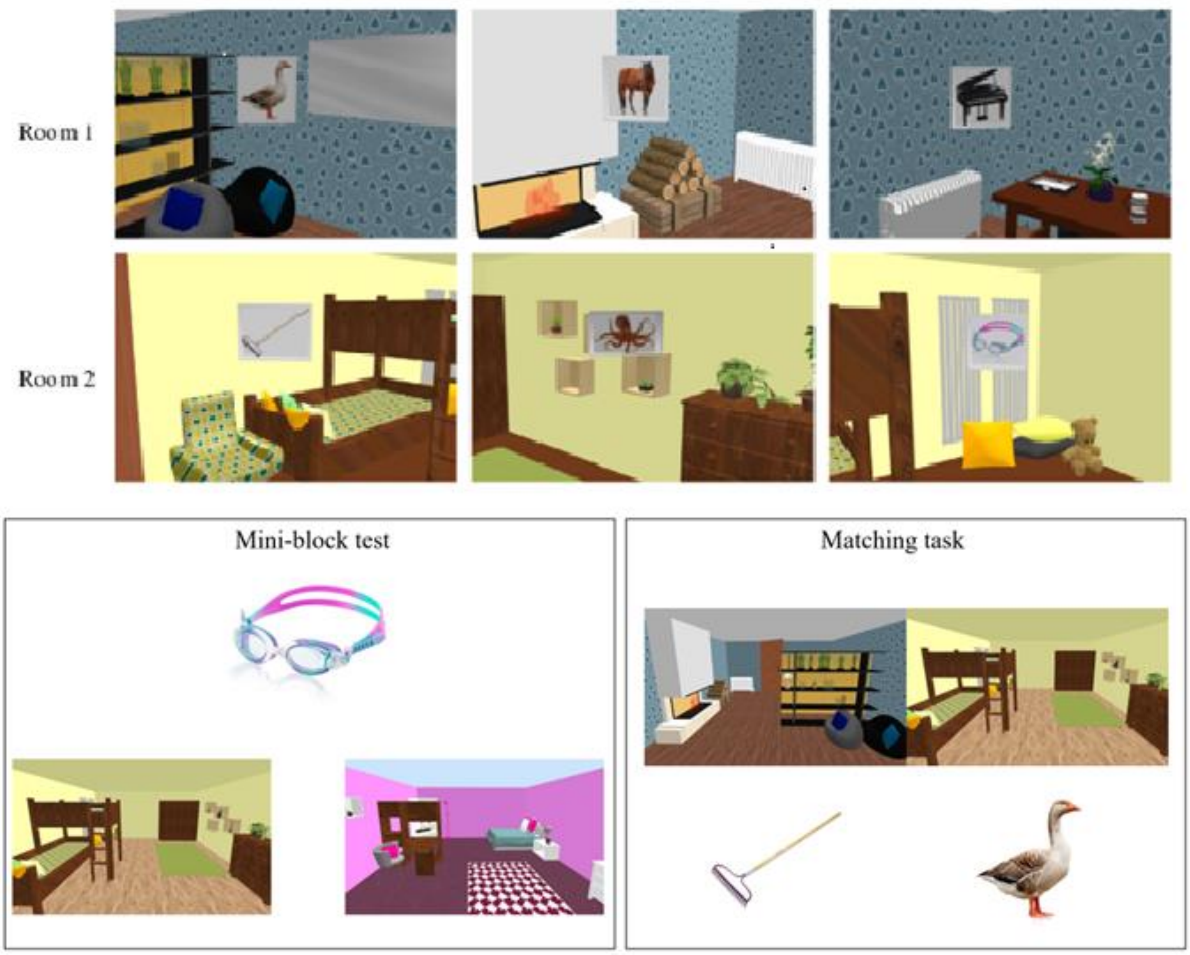


*Figure S1. Top panel:* An example of what was seen during the training videos. This is a 'Mixed Category Building' that contains items from different semantic categories. An example walk-through video of this building can be watched following this link: https://www.youtube.com/watch?v=XVHhOh3BF74&feature=youtu.be. *Bottom panel:* Example of training tests. On the left is an example of the mini-block test depicting the probe item at the top and two room screenshots below. The target screenshot is the room the object was presented in at training (left). The distractor screenshot (right) is from a different building, which the object did not belong to. On the right is an example of the matching task depicting two rooms from the same building and two items that belong to each room (rake-right room, goose-left room) which participants needed to match.

Study 2: Materials and Results

Figure S2 shows example stimuli used in Study 2. Our localisers focussed on the Scenes and Objects conditions.

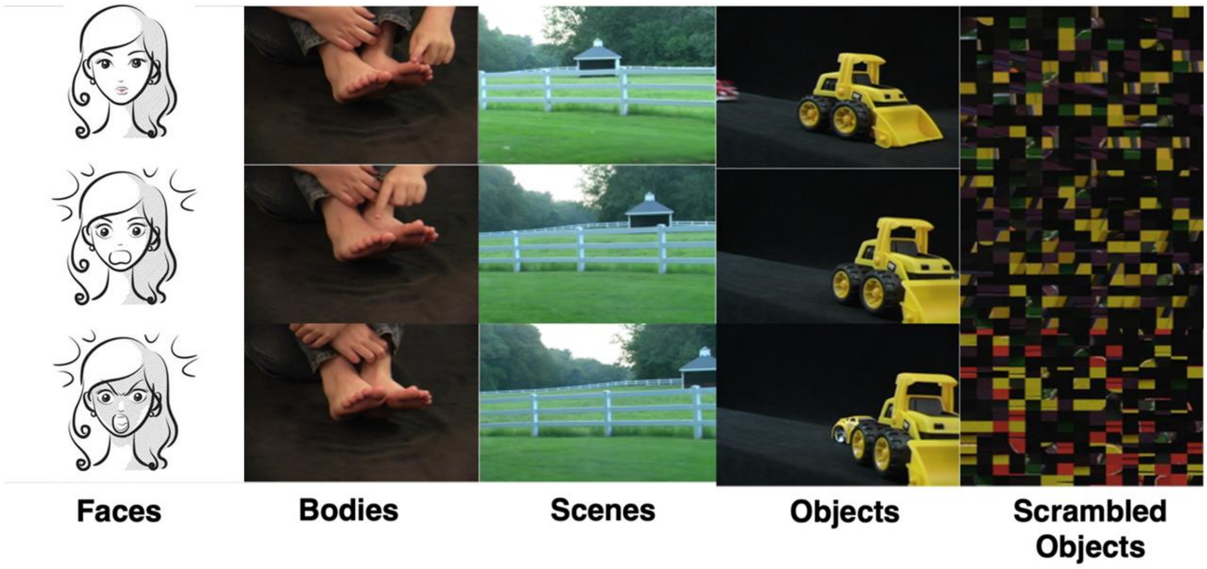


*Figure S2.* Examples of the dynamic images taken from the 3-s movie clips depicting faces, bodies, scenes, objects, and scrambled objects. Still images taken from the beginning, middle and end of the corresponding movie clip. The stimuli corresponding to the 'Faces' condition were changed to line drawings to make this material suitable for hosting in bioRxiv preprint server. The actual stimuli shown can be consulted in the OSF collection associated with this paper (https://osf.io/sh79m/)

As shown in the left panel of Figure S3, the objects over scrambled objects contrast revealed activation in areas associated with the processing of objects and grasping, such as bilateral LOC, fusiform, parietal and precuneal cortex. The scenes over objects contrast showed activation in medial occipital and retrosplenial cortex, associated with scene processing. Since we were interested in using these activation maps as masks to determine relevant regions of the visual end of our pathways, we masked these effect maps by large-scale subnetworks implicated in vision from an influential parcellation (Yeo et al., 2011; visual central and visual peripheral networks combined). The resulting masks showed some voxels in common, and since the aim of this analysis was to identify areas that preferentially respond to objects or scenes, we excluded these voxels in a final step (i.e., we removed all voxels from the objects mask that were also part of the scenes mask, and vice versa). These results can be consulted on the right panel of Figure S3.

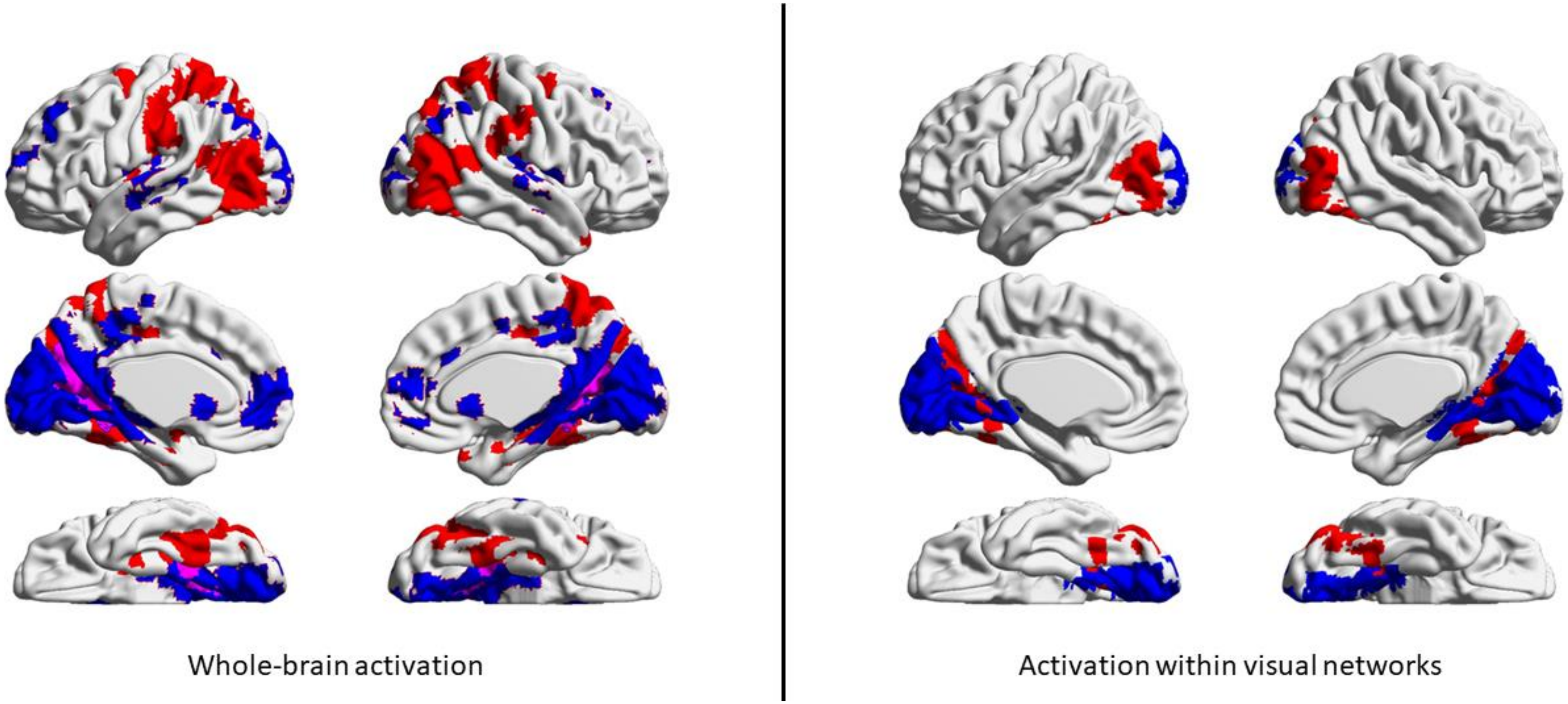


*Figure S3. Left panel:* Areas associated with the passive viewing of objects are shown in red, and those associated with the passive viewing of scenes are shown in blue; areas that responded to both objects and scenes are shown in purple. *Right panel:* The results of the localiser shown in the left panel were further constrained to contain only voxels that overlapped with the visual networks in Yeo's 17 network parcellation. Common voxels (the ones that responded to both objects and scenes) have been removed.

Supplementary Analysis: Resting-state maps with stricter thresholding

Since the pattern of results obtained from resting-state analysis can be sensitive to threshold decisions, we reproduced the group-level maps depicted in Figures 2E and 3E using a stricter threshold. We originally thresholded these results using the default of the CONN software package (cluster-forming threshold of $p=.05$, equivalent to $T=1.65$). For increased rigour, we reproduced the thresholded maps from Figures 2E and 3E further increasing the threshold from $p=.05$, equivalent to $T=1.65$, to $p=.001$, equivalent to $T=3.1$. The resulting maps were very similar, showing minimal change, with a spatial correlation of $r > .99$ between the strict and lax threshold versions of the maps for both the probe and decision seeds. These maps can be downloaded from the OSF collection associated with this project.

Supplementary Analysis: Eroded Masks Replication Analysis

The proximity of visual peripheral and DMN-C network borders is a property of the organisation of these networks (Silson et al., 2019; Steel et al., 2021). However, this could give rise to the potential for spatial mixing of the resting-state signal during intrinsic connectivity analysis due to smoothing, falsely inflating the strength of connectivity, since our visual and DMN masks for the spatial context task showed spatial adjacency. To address this concern, we re-analysed the resting-state data presented in panels b and c of Figure 4 (connectivity from DMN decision regions to visual probe regions and vice versa) by eroding the visual probe and DMN decision ROIs for the spatial context task using fslmaths. We eroded the masks until the smallest gap between them exceeded the size of our 6mm FWHM smoothing kernel, which eliminates the potential for spatial mixing of signals due to ROI adjacency. The eroded ROIs can be consulted in the OSF collection associated with this project. We repeated the ANOVAs associated with these analyses. The results, presented in Supplementary Figure S4 below, confirmed the pattern of findings reported in the main analysis. We did not erode the respective ROIs for the semantic task, given that adjacency is not an issue for the ROIs derived from that task.

The Visual-to-DMN ANOVA showed main effects of seed ($F(1,190)=22.82$, $p<.001$), ROI ($F(1,190)=9.48$, $p=.002$) and a seed by ROI interaction ($F(1,190)=67.02$, $p<.001$). Post-hoc contrasts confirmed there was stronger connectivity between object probe regions and semantic versus spatial context decision regions ($t(190)=3.38$, $p<.001$), and between scene probe regions and spatial context versus semantic decision regions ($t(190)=-7.66$, $p<.001$).

The DMN-to-Visual ANOVA confirmed this pattern: again, there was a main effect of ROI (F(1,190)=4.3, p=.039) and a seed by ROI interaction (F(1,190)=57.59, p<.001), with post-hoc contrasts confirming stronger intrinsic connectivity between DMN regions implicated in sematic decisions and object probe regions (t(190)=5.06, p<.001), and between DMN regions engaged by spatial context decisions and scene probe regions (t(190)=3.25, p=.001).

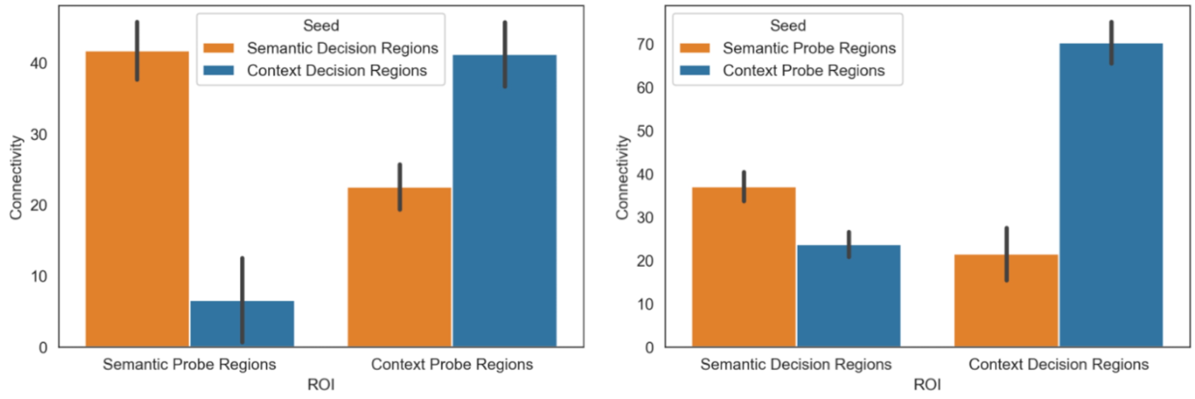


Supplementary Figure S4. Results of the re-analysis of intrinsic connectivity between semantic and context visual probe and DMN decision regions.

<u>Supplementary Analysis: Replicating resting-state connectivity pathways with task-independent ROIs</u>

We performed a supplementary analysis using task-independent ROIs to confirm that the intrinsic connectivity-based pathways could be identified even when the seeds and ROIs were not derived from the same task. In this analysis, we used the same seeds as the main analysis (Figure 4a; the conjunction of semantic and context decision activation within DMN and object and scene probes within visual networks), while the ROIs were either visual localiser masks for objects and scenes, or DMN subsystems from the Yeo et al. (2011) parcellation. The results can be consulted in Figure S5. The DMN-to-visual ANOVA revealed main effects of DMN seed (F(1,190)=137.72, p<.001), visual ROI based on the localisers from Study 2 (F(1,190)=23.87, p<.001) and their interaction (F(1,190)=20.92, p<.001). Post-hoc tests revealed that DMN regions associated with spatial context decisions showed stronger connectivity to both visual regions associated with viewing scenes, and visual regions associated with viewing objects, relative to DMN regions associated with semantic decisions (scenes: t(190)=11.57, p<001; objects: t(190)=7.35, p<001). The significant interaction term reveals that this difference between context and semantic seeds was more pronounced for the scene than the object localiser regions, confirming a greater importance of the context pathway for making decisions based on visual scene information.

The visual-to-DMN ANOVA revealed main effects of visual seed (F(1,190)=33.98, p<.001), DMN ROI (F(1,190)=65.46, p<.001) and their interaction (F(2,380)=119.14, p<.001) . Post-hoc tests revealed that visual regions associated with viewing object probes showed stronger connectivity to FT-DMN regions relative to regions associated with viewing spatial context probes (t(190)=3.22, p=002), while regions associated with viewing spatial context probes showed stronger connectivity to core and MT-DMN regions than regions associated with viewing object probes (Core DMN: t(190)=4.41, p<.001; MT-DMN: t(190)=11.7, p<.001).

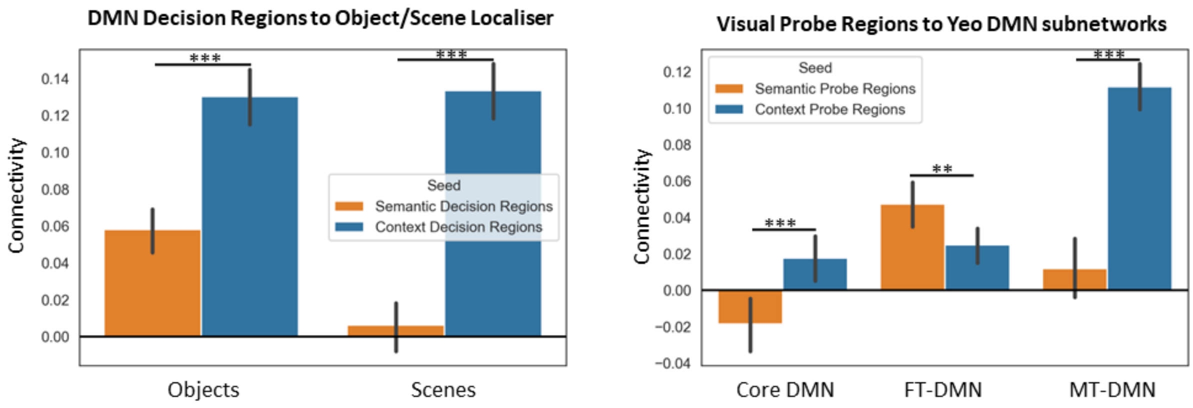


*Figure S5.* Connectivity of the DMN decision regions to the object/scene localiser from Study 2, and that of the visual probe regions to FT, MT and core DMN of Yeo's 17-network parcellation.

Supplementary Analysis: Replicating pathways' structural connectivity from the DMN end

To confirm dissociable pathways based on structural connectivity can be identified not only when examining visual-to-DMN regions, but also the reverse, we performed a supplementary analysis seeding the DMN, and examined the strength of connections to visual ROIs (defined using activation to object and scene probes in Study 1 masked by visual networks). The results can be consulted on Figure S6. The DMN-to-visual ANOVA showed a main effect of visual ROI (F(1,163)=506.5, p<.001) and a seed by ROI interaction (F(1,163)=215.27, p<.001). Post-hoc tests revealed that both DMN seeds, associated with semantic and spatial context decisions, showed stronger connectivity to visual regions responding more to scenes than to objects. However, this connectivity difference was greater for DMN regions activated by spatial context than semantic decisions (semantic DMN: t(163)=3.92, p<.001; spatial context DMN: t(163)=2381.12, p<.001).

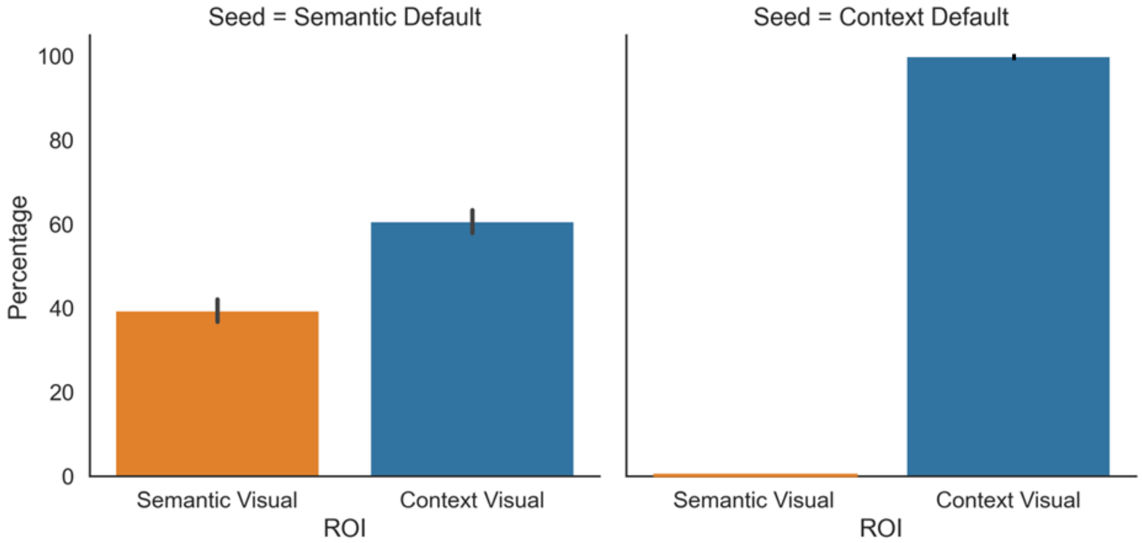


Figure S6. The y axis of the bar plots shows the percentage of streamlines from each DMN seed that terminate in each visual ROI (shown in the x axis). Seeds and ROIs can be consulted in Figure 4a. The error bars depict the standard error of the mean.

### Supplementary Analysis: Effects of task demands on pathway connectivity

*Methods*

In order to test for distinct semantic and spatial memory pathways that connect visual regions to distinct subnetworks of the DMN, we conducted a psychophysiological interaction (PPI) analysis. Semantic and spatial context visual seeds were created from the univariate activation to object and scene probes in the semantic and spatial tasks respectively, masked by Yeo et al. (2011) 7-network parcellation visual network. The timeseries of these seeds were then extracted after pre-processing. We then ran two separate models (one for each seed), which examined the main effect of the experimental condition (i.e., *SCB trials of the semantic task, MCB trials of the semantic task, SCB trials of the spatial context task and MCB trials of the spatial context task*). These models included all eight regressors from the basic task model of Study 1 described in section 2.1.4., a PPI term for each of the seven conditions and phases of the task (SCB/MCB trials of the probe, dots and decision phases, and the arrow task), as well as the time series of the visual probe seeds, using the generalized psychophysiological interaction (gPPI) approach (McLaren et al., 2012). The regressors were not orthogonalized. All runs of each task were combined using fixed-effects analyses for each participant, which allowed us to extract the connectivity parameters for each experimental condition for each participant in each seed model.

*Results and Discussion*

We examined how connectivity within the pathways changes depending on task demands in a Psychophysiological Interaction (PPI) analysis. We took the visual regions showing differential activation to object and scene probes as seeds (shown in Figures 2e and 4a), while the ROIs were regions sensitive to semantic and spatial context decisions within the DMN (shown in Figures 3e and 4a). We anticipated that the scene probe regions would increase their connectivity to spatial context decision regions during the decision phase of the spatial context task, whilst the object probe regions would increase their connectivity to semantic decision regions during the decision phase of the semantic task. A repeated-measures ANOVA including task (semantic/spatial context), seed (object/scene probe), ROI (semantic/spatial context decision) and condition (SCB/MCB) as factors revealed two-way interactions for task by seed ($F(1,26)=10.85, p=.003$) and seed by ROI ($F(1,26)=8.57, p=.007$), as well as a three-way interaction for task by seed by ROI ($F(1,26)=5.2, p=.031$). Since we found no effect of condition, we averaged across this factor for the following analyses. The results are shown in Supplementary Figure S7. To understand the three-way interaction, separate two-way ANOVAs using seed (object/scene probe regions) and ROI (semantic/spatial context decision regions) as factors were computed for the spatial context and semantic tasks. The semantic task showed a main effect of seed ($F(1,26)=6.97$, $p=.014$), but no effect of ROI or interaction: the object seed was more connected to both semantic and spatial context DMN decision regions during the semantic task. The spatial context task showed a main effect of seed ($F(1,26)=5.89$, $p=.022$), and a seed by ROI interaction ($F(1,26)=10.25$, $p=.004$). Post-hoc t-tests showed that the scene probe regions were more connected to spatial context decision regions during the spatial context task than object probe regions ($t(26)=3.52$, $p=.002$). In contrast, there was no difference in connectivity between these two seeds and the semantic decision regions.

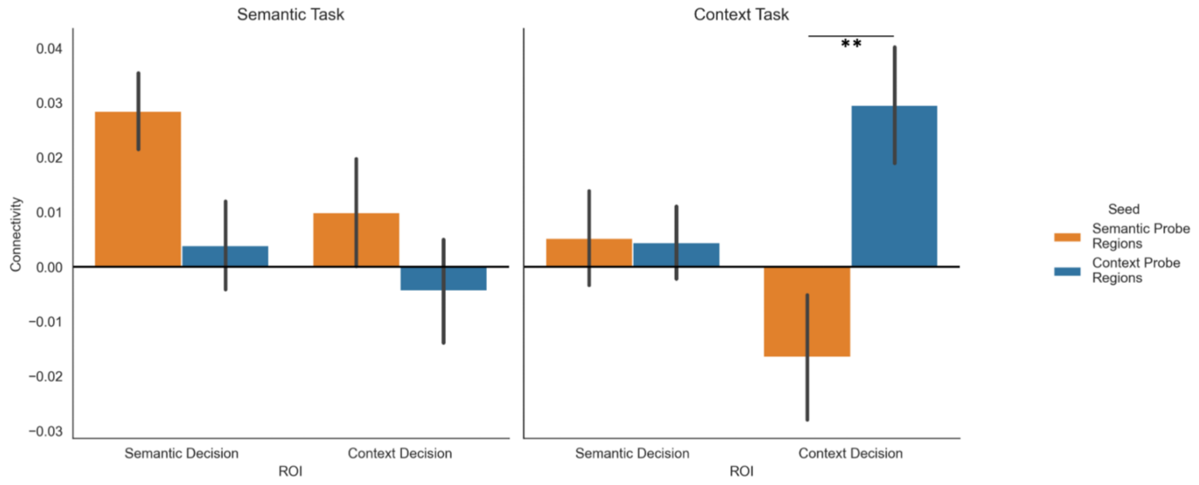


Figure S7. Psychophysiological interaction analysis of the connectivity from the spatial context and semantic probe regions to FT and MT-DMN subnetworks. This analysis collapses the SCB and MCB conditions, which showed no significant differences. Note. * = p<.05, ** = p<.01.

In sum, the psychophysiological interaction models characterised how inputs to the pathways are flexibly configured to suit our current goals. The visual ends of the pathways showed opposing patterns of connectivity to spatial context DMN regions depending on the task.

### Supplementary Analysis: Individual Location of pathways in whole-brain gradients

Our analysis of the location of the pathways in whole-brain gradient connectivity space showed that peak responses during semantic decisions occurred in more abstract, less visual regions of the DMN relative to spatial context decisions. However, the scatterplots in the top panel of Figure 5 do not allow to distinguish whether these effects took place at the individual level, since the data points are not linked across tasks. In light of this, we plotted the same data comparing the gradient values for the peak responses in each of our tasks at the participant level. The peaks for each participant across tasks are linked with a line. Cases where the pattern was reversed are highlighted with dashed lines (7/27 participants in each gradient, see Supplementary Figure S8). This analysis showed that in the majority of cases, at the individual level, the pattern of group-level results held.

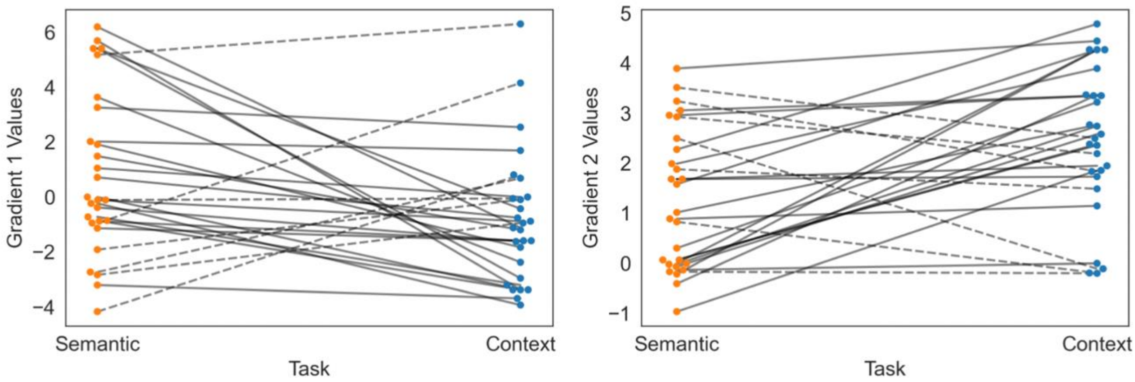


Figure S8. Location in the two principal gradients of the peak response per participant for semantic and spatial context decisions. Dashed lines highlight the cases that contradict the pattern found in the group level analysis.

Supplementary Analysis: Multivariate Response to Same- versus Mixed-Category Buildings

*Methods*

Since distinct but adjacent regions were associated with semantic and spatial context decisions, we asked what they represented during probe presentation in MCB and SCB trials using Representational Similarity Analysis (RSA). Probes were presented across decision and no-decision trials, allowing a large number of probe responses to be included in the analysis. We examined the voxels that responded to contrasts between semantic and spatial context decisions (including all significant, suprathreshold voxels at the group level within a single region-of-interest). We constructed semantic similarity matrices for each participant using all pairs of trials from the semantic task, encoding category similarity on a scale of 0 to 2. Pairs of trials that shared a specific category (e.g., birds) were assigned the strongest value (2), while those that shared only their superordinate category (animals versus man-made objects) were assigned the middle value (1); pairs of trials from different superordinate categories were assigned the weakest value (0). We also constructed spatial context similarity matrices for each participant, encoding the relationships between rooms and buildings on a scale of 0 to 2. Pairs of trials belonging to the same room were assigned the strongest value (2), while those belonging to different rooms of the same building were assigned a middle value (1); trials that belonged to different buildings were assigned the lowest value (0).

*Single-trial Estimation:* GLMs were performed separately to estimate the activation pattern for each of 144 trials during the probe phase in the two tasks. A Least Square–Single (LSS) approach was used, in which the trial of interest was modelled as one regressor, with all other trials modelled as separate regressors (Mumford et al., 2012). These models included eight regressors: (1) the probe phase of interest (SCB or MCB); (2 and 3) all other probe phases (SCB and MCB); (4) Dots SCB; (5) Dots MCB; (6) Decision SCB; (7) Decision MCB; (8) Arrow trials. Since the analysis focused on probe presentation rather than decisions, incorrect trials were not excluded. Each event was modelled at the time of stimulus onset and convolved with a canonical hemodynamic response function (double gamma), whereas the fixations were treated as an implicit baseline. Pre-whitening was applied. The same pre-processing procedure as in the univariate analysis was used except that no spatial smoothing was applied. This voxel-wise GLM was used to compute the activation associated with each trial, using the t-map for Representational Similarity Analysis to increase reliability by normalizing for noise (Walther et al., 2016).

*Second-Order Representational Similarity Analysis:* A searchlight approach compared semantic and spatial context similarity matrices with neural similarity matrices. Neural pattern similarity was estimated for cubic regions of interest (ROIs) within t-maps for each trial, containing 125 voxels surrounding a central voxel (Fairhall and Caramazza, 2013; Gao et al., 2022; Leshinskaya et al., 2017; Malone et al., 2016; Stolier and Freeman, 2016; Viganò and Piazza, 2020; Wang et al., 2017). In each of these cubes, we derived a neural similarity matrix from Pearson correlations of pairs of trials. We excluded any pairs presented in the same run to avoid any autocorrelation. Spearman's rank correlation was used to measure the alignment between task and brain-based models during the probe phase. The resulting coefficients were Fisher's z transformed and then entered into a group level analysis carried out using FSL's Randomise (Anderson and Robinson, 2001; Winkler et al., 2014) (5,000 permutations with Threshold-Free Cluster Enhancement), thresholding the results at $p < .05$.

We also performed cross-task similarity analysis, correlating semantic similarity to the neural similarity matrix from the spatial context task (and vice versa). If participants use semantic information learned during training to guide spatial context decisions, or spatial context information from training to facilitate semantic decisions, we might be able to identify regions sensitive to semantic and spatial context information across tasks. This should only be the case in SCB and not MCB trials.

*Results and Discussion*

The univariate analysis in the main text shows that when there is no alignment between spatial context and semantic information (in MCB trials), the heteromodal areas that are activated by the task show higher pathway-specific connectivity. In contrast, when information integration across space and meaning is facilitated by the structure of the task, spatial context trials show more activation in regions with lower connectivity to the spatial context pathway, but higher connectivity to the semantic pathway. In this way, right angular gyrus was found to have a potential role in integrating the visual-to-DMN pathways.

A follow-up analysis used a multivariate approach to establish how neural patterns related to the task reflected information integration. We performed Representational Similarity Analysis (RSA) using a searchlight approach within a mask that combined semantic and spatial context task decision maps, using data acquired during the probe phase (since there were more probe than decision time-points). This method allowed us to select regions sensitive to semantic and spatial context information, while ensuring that the search space was not derived from the same data used for the RSA analysis. First, we asked if we could detect regions sensitive to category during the semantic task and sensitive to location during the spatial context task, in the MCB trials. There were regions that represented semantic and spatial context similarity in bilateral and left LOC respectively (Figure S9, panel a). Next, we performed a cross-task representational similarity analysis in the SCB trials to identify areas that represented information relevant to one task in the other (e.g., areas that represented semantic information during the spatial context task and vice versa). The results of this analysis revealed right LOC regions that captured spatial context information during the semantic task (Figure S9, panel b). No medial regions were found in these analyses.

Finally, we investigated the intrinsic connectivity of these multivariate clusters to the semantic and spatial context pathways (Figure 4d and Figure S9, panel c), to establish whether cross-task RSA regions thought to support integration have an intermediate pattern of connectivity to both pathways. We used the MCB semantic and spatial context RSA clusters and the cross-task RSA result from Figure S9a-c as seeds in a seed-to-ROI analysis of intrinsic connectivity using independent data from Study 3. We performed a 2 two-way repeated measures ANOVA, using a 3 x 2 design, entering seed (semantic, spatial context and cross-similarity RSA results), and pathway (ROIs in Figure S9, panel c) as factors. The results can be seen in Figure S9 panel d. There were main effects of seed ($F(1.54,293.32)=194.24$, $p<.001$), ROI ($F(1,190)=290.07$, $p<.001$) and their interaction ($F(1.58,300.36)=123.36$, $p<.001$). Post-hoc comparisons confirmed that the spatial context pathway was equally connected to the spatial context RSA cluster and to the cross-task RSA cluster (spatial context > cross-task: $t(190)=-.155$, $p>.05$), with both of these clusters being significantly more connected than the semantic RSA cluster (spatial context > semantic: $t(190)=3.34$, $p=.002$; cross-task > semantic: $t(190)=4.41$, $p<.001$). The semantic pathway was most connected to the semantic RSA cluster, less connected to the cross-task RSA cluster, and least connected to the spatial context RSA cluster (semantic > cross-task: $t(190)=9.2$, $p<.001$; cross-task > spatial context: $t(190)=16.31$, $p<.001$). In this way, the cross-task representation of spatial context information in visual regions during the semantic task showed an intermediate pattern of connectivity (particularly to the semantic pathway).

These cross-talk regions, in right lateral occipital cortex, have been implicated in the integration of objects with their spatial location, allowing object coherence in space in the face of saccadic movements that occur in natural vision while navigating environments (McKyton and Zohary, 2007). Another fMRI study investigating the structure of identity-related and location-related representations in visual regions found an interaction effect of these aspects of knowledge for objects positioned in expected spatial locations, in a similar fashion to our study (Gronau et al., 2008).

For completion, we present the results of RSA analysis of SCB trials in Figure S10 below.

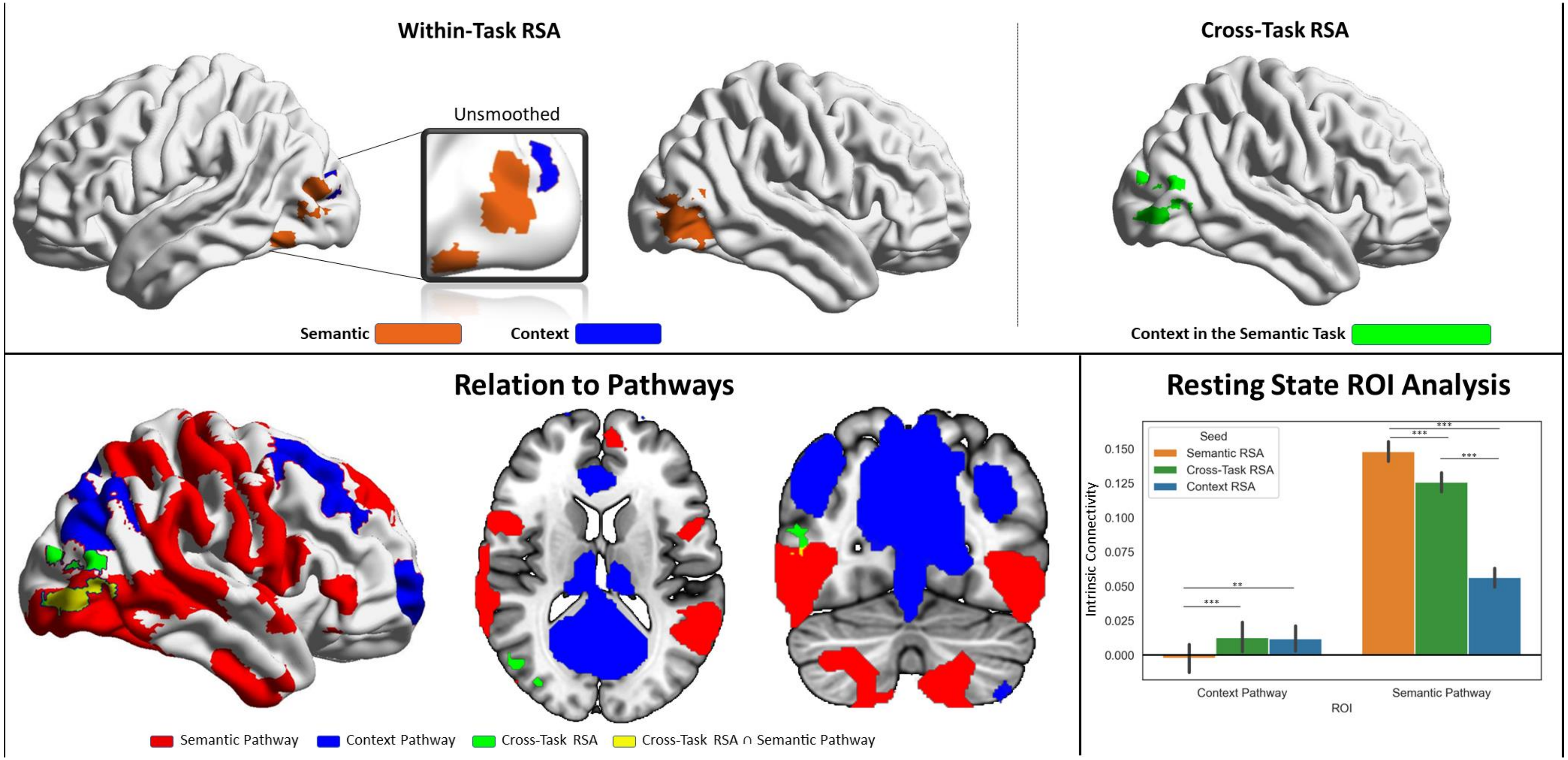


*Figure S9.* Results of the representational similarity analysis. Top left panel: within-task RSA results correlating BOLD activity from the probe phase of semantic mixed-category building trials with the semantic similarity matrix described in the Methods section, and BOLD activity from the probe phase of context mixed-category building trials with the context similarity matrix. Top right panel: cross-task similarity analysis correlating BOLD activity from semantic trials with the context similarity matrix. Bottom panel: The left part depicts the spatial relations of the cross-task similarity analysis cluster with the semantic and context pathways outlined in Figure 4; the right part shows Intrinsic connectivity seed-to-ROI results using the within- and cross-task RSA clusters shown in the top panel as seeds and the pathways as ROIs. The error bars depict the standard error of the mean. *** $p < .001$, ** $p < .01$.

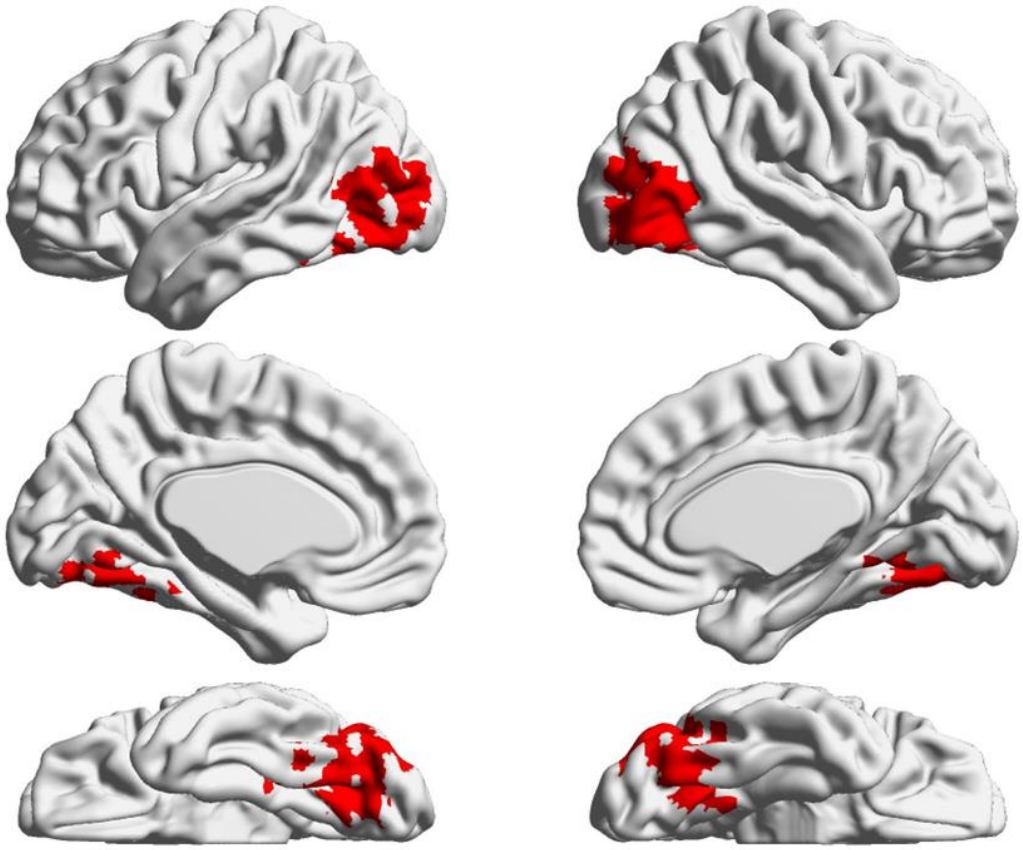

*Figure S10*. Representational Similarity Analysis results for the Same-Category Building trials of the Probe phase of the Semantic task. No significant voxels were identified for the spatial context task.

Supplementary Table S1. Cluster Information for Neuroimaging Results from Study 1

| | | | | Coordinates (in mm) | | | |
|---|---|---|---|---|---|---|---|
| Analysis | Hemisphere | Cluster Peak | Z | x | y | z | Cluster Volume |
| Probe: Semantic > Spatial Context | Left | Lateral Occipital Cortex, Inferior Division | 4.78 | -50 | -74 | 4 | 932 |
| | Right | Lateral Occipital Cortex, Inferior Division | 4.58 | 50 | -68 | 0 | 534 |
| | Left | Parietal Operculum Cortex | 3.9 | -54 | -34 | 22 | 218 |
| Probe: Spatial Context > Semantic | Right | Occipital Pole | 6.64 | 14 | -92 | 2 | 9931 |
| | - | Paracingulate Gyrus | 3.82 | 0 | 16 | 52 | 495 |
| | Left | Frontal Pole | 3.78 | -30 | 60 | 14 | 342 |
| | Left | Frontal Orbital / Insular Cortex | 4.42 | -30 | 26 | -2 | 236 |
| | Right | Middle Frontal Gyrus | 3.73 | 32 | 2 | 64 | 218 |
| | Left | Lateral Occipital Cortex, Superior division | 3.7 | -36 | -60 | 54 | 218 |
| Decision: Semantic > Spatial Context | Left | Lateral Occipital Cortex, Inferior Division | 5.24 | -50 | -74 | 4 | 2438 |
| | Right | Lateral Occipital Cortex, Inferior Division | 5.32 | 50 | -70 | 2 | 1605 |
| | Left | Middle / Inferior Frontal Gyrus | 4.6 | -50 | 28 | 24 | 1369 |
| | Left | Middle Frontal / Precentral Gyrus | 4.57 | -52 | 8 | 40 | 385 |
| | Left | Intracalcarine Cortex | 3.92 | -6 | -76 | 12 | 363 |
| | Left | Frontal Pole / Superior Frontal Gyrus | 3.55 | -12 | 40 | 52 | 217 |
| | Right | Frontal Operculum Cortex | 4.04 | 46 | 18 | -6 | 207 |
| Decision: Spatial Context > Semantic | Right | Occipital Pole | 4.62 | 4 | -94 | -6 | 1543 |
| | Right | Temporal Occipital Fusiform Cortex | 5.34 | 30 | -46 | -8 | 420 |
| | Right | Precuneous Cortex | 4.17 | 20 | -58 | 18 | 253 |
| | Left | Lingual Gyrus | 5.12 | -24 | -46 | -8 | 251 |
| Probe: Spatial Context Mixed > Same | Right | Lateral Occipital Cortex, Superior division | 4.06 | 50 | -70 | 32 | 294 |
| | Left | Precuneous Cortex | 3.71 | -8 | -70 | 28 | 232 |
| Probe: Spatial Context Same > Mixed | Right | Angular Gyrus | 3.85 | 58 | -50 | 44 | 222 |
| Probe: Semantic Mixed > Same | Left | Lateral Occipital Cortex, Superior division | 3.99 | -36 | -60 | 54 | 610 |
| | Right | Superior Parietal Lobule | 3.51 | 38 | -54 | 58 | 364 |
| | Left | Lateral Occipital Cortex, inferior division | 3.36 | -50 | -66 | -18 | 286 |
| | Right | Inferior Temporal Gyrus, temporooccipital part | 3.45 | 50 | -60 | -22 | 278 |
| | Right | Lateral Occipital Cortex, Superior division | 3.57 | 40 | -78 | 38 | 265 |

Supplementary Table S2. Paired t-tests contrasting spatial similarity of participant-level activation with group-level context and semantic pathways and non-pathways

| Individual Task Responses | Group-defined pathways | Non-pathway conjunctions | t | df | Two-sided p |
|---|---|---|---|---|---|
| Context Decision | Context Pathway (Context Probe and Context Decision) | Context Probe and Semantic Decision | 76.25 | 190 | 2E-144 |
| Context Decision | Context Pathway (Context Probe and Context Decision) | Semantic Probe and Context Decision | 70.89 | 190 | 1.3E-138 |
| Semantic Decision | Semantic Pathway (Semantic Probe and Semantic Decision) | Context Probe and Semantic Decision | 17.55 | 190 | 1.22E-41 |
| Semantic Decision | Semantic Pathway (Semantic Probe and Semantic Decision) | Semantic Probe and Context Decision | 52.95 | 190 | 1E-115 |
| Context Probe | Context Pathway (Context Probe and Context Decision) | Context Probe and Semantic Decision | 2.36 | 190 | 0.019292 |
| Context Probe | Context Pathway (Context Probe and Context Decision) | Semantic Probe and Context Decision | 31.11 | 190 | 1.74E-76 |
| Semantic Probe | Semantic Pathway (Semantic Probe and Semantic Decision) | Context Probe and Semantic Decision | 14.2 | 190 | 1.14E-31 |
| Semantic Probe | Semantic Pathway (Semantic Probe and Semantic Decision) | Semantic Probe and Context Decision | 18.01 | 190 | 5.68E-43 |

Note. T values represent contrasts between the correlations of individual task responses and group-defined pathways versus these same responses correlated with non-pathway conjunctions (pathway > non-pathway)
